# Changes in high-frequency aperiodic 1/f slope and periodic activity reflect post-stimulus functional inhibition in the visual cortex

**DOI:** 10.1101/2023.10.01.560381

**Authors:** Viktoriya O. Manyukhina, Andrey O. Prokofyev, Tatiana S. Obukhova, Tatiana A. Stroganova, Elena V. Orekhova

## Abstract

It has been shown that cessation of intensive sensory stimulation is associated with a transient increase in functional inhibition in the sensory cortical areas. However, the electrophysiological correlates of this post-stimulus inhibition in the human brain have not been properly investigated.

To investigate post-stimulus inhibition, we recorded magnetoencephalogram (MEG) at rest and after cessation of visual stimulation of varying intensity (high-contrast gratings drifting at a slow, medium, or high rate) in 25 healthy women aged 18-40 years. We analyzed condition- and intensity-related changes in MEG parameters sensitive to functional inhibition: periodic alpha-beta power, peak alpha frequency (PAF), and 1/f aperiodic slope. We also investigated the association of these parameters with sensory sensitivity and avoidance assessed by a questionnaire. To evaluate the influence of hormonal status on the studied parameters, participants were examined twice, during the follicular and luteal phases of the menstrual cycle (MC).

Regardless of MC phase, increasing intensity of the preceding visual stimulation resulted in increased posterior post-stimulus alpha-beta power, PAF, and a more negative/steeper aperiodic (1/f) slope of the power spectrum in the 35-45 Hz range. Compared to rest, the post-stimulus period was characterized by higher PAF, a steeper 35-45 Hz 1/f slope in posterior cortical areas, and a widespread increase in beta power. While condition- and intensity-dependent changes in alpha-beta power and 1/f slope were correlated, changes in PAF did not correlate with either of them. Greater intensity-dependent increase in visual alpha-beta power predicted higher subjective sensory sensitivity/avoidance, suggesting stronger regulatory top-down modulation of visual cortex in individuals with heightened sensitivity.

Our results show that several MEG parameters concordantly indicate a post-stimulus enhancement of inhibition that is proportional to the intensity of the preceding visual stimulation. While post-stimulus changes in alpha-beta power and 1/f slope may share some common mechanisms, changes in PAF reflect a distinct aspect of inhibitory regulation. Our results inform potential inhibition-based biomarkers for clinical and translational research.

**Key points:**

- The cessation of visual stimulation leads to a transient increase in functional inhibition.
- Changes in MEG parameters reflect post-stimulus enhancement of inhibition.
- These changes scale with the intensity of the preceding stimulation.

## Introduction

A balance between neural excitation and inhibition (E/I balance) in neural circuits is crucial for the normal functioning of the brain and its disruption is associated with a variety of neuropsychiatric disorders (Anticevic & Murray, 2017; Sohal & Rubenstein, 2019). Recent advances in understanding the role of E/I balance in the etiology of these disorders instigated a search for its non-invasive and accessible biomarkers that could help to stratify patients according to the dominant neural deficit and be used in clinical and translational research.

Several classes of non-invasive electrophysiological biomarkers of the E/I balance have been suggested which are usually based on electroencephalographic (EEG) and magnetoencephalographic (MEG) parameters estimated ‘at rest’ or during sensory stimulation (see (Ahmad et al., 2022) for review). Yet, the neural activity recorded *after cessation* of intensive sensory stimulation may provide additional valuable information regarding the link between MEG/EEG parameters and regulation of the E/I balance. According to cellular (Shmuel, Augath, Oeltermann, & Logothetis, 2006; Q. Zhang et al., 2023), blood-oxygen-level dependent (BOLD) functional magnetic resonance imaging (fMRI) (Havlicek et al., 2015; Mullinger, Mayhew, Bagshaw, Bowtell, & Francis, 2013; Sadaghiani, Uğurbil, & Uludağ, 2009), MEG (Stevenson, Brookes, & Morris, 2011), and EEG (Mullinger, Cherukara, Buxton, Francis, & Mayhew, 2017; Mullinger et al., 2013) studies, the period after stimulation is dominated by inhibition, i.e., suppression of excitatory neuronal activity. Since the post-stimulus neural suppression may originate from multiple sources, following Mayhew et al (Mayhew, Coleman, Mullinger, & Can, 2022) and Iemi et al (Iemi et al., 2022) here we will use the term post-stimulus inhibition (or functional inhibition) to refer to a decrease in the E/I ratio that can occur via activation of GABAergic inhibitory interneurons and/or via decreased excitatory drive (e.g., downregulation of norepinephrine and acetylcholine).

In the primate visual cortex, termination of visual input is immediately followed by local decreases of multiunit neural activity below the baseline level (Logothetis, Pauls, Augath, Trinath, & Oeltermann, 2001; Shmuel et al., 2006). In human fMRI, a cessation of visual stimulation is accompanied by the ‘undershoot’ - a negative (i.e. below pre-stimulus baseline) BOLD response that at least in part is accounted for by the shift of the E/I ratio to inhibition (Havlicek et al., 2015; Mullinger et al., 2013; Sadaghiani et al., 2009). Recent studies have shown that fMRI BOLD undershoot is reduced in elderly people (Mayhew et al., 2022) and in people with some neuropsychiatric disorders (Havlicek et al., 2015; Murray, Kolodny, Schallmo, Gerdts, & Bernier, 2020), suggesting weakened neural inhibition and altered E/I ratio.

Whereas the hemodynamic undershoot in humans has been extensively investigated (J. J. Chen & Pike, 2009; Haigh, Cooper, & Wilkins, 2015; Havlicek et al., 2015; Mullinger et al., 2017; Mullinger et al., 2013; Sadaghiani et al., 2009; Stevenson et al., 2011), less attention has been paid to the MEG/EEG indices of post-stimulus inhibition. Since MEG and EEG directly reflect neural functioning, they may appear even more informative than the BOLD fluctuations regarding the regulation of E/I balance in healthy and diseased human brain.

In EEG and MEG, the cessation of visual stimulation is followed by an increase in alpha and beta power in the visual cortex (Mullinger et al., 2017; Stevenson et al., 2011). Similarly to fMRI BOLD undershoot, the post-stimulus alpha-beta synchronization differs from that observed during baseline, and in this case, it *exceeds* the pre-stimulus baseline level (Mullinger et al., 2017; Stevenson et al., 2011). Importantly, this post-stimulus alpha ‘overshoot’ correlates with the fMRI undershoot on a trial-by-trial basis (Mullinger et al., 2017), which suggests that both phenomena reflect inhibition associated with the termination of visual stimulation. Indeed, increases in periodic alpha and beta power have been previously associated with reduced excitability as evidenced by their negative correlations with high-frequency broadband gamma activity (Iemi et al., 2022) and neuronal spiking (Chapeton, Haque, Wittig, Inati, & Zaghloul, 2019; Dougherty, Cox, Ninomiya, Leopold, & Maier, 2017; Haegens, Nácher, Luna, Romo, & Jensen, 2011).

The frequency of alpha oscillations, although not investigated in the context of post-stimulus inhibition, has also been shown to reflect state-related changes in cortical excitability. For example, the increased frequency of posterior alpha oscillations during the retention period of a memory task is thought to reflect inhibition of sensory processing, because it is accompanied by slower reaction time and decreased neural responses to external stimuli (Babu Henry Samuel, Wang, Hu, & Ding, 2018). Furthermore, the spontaneous fluctuations of alpha frequency are inversely associated with BOLD activity in the visual cortex (Babu Henry Samuel et al., 2018). Given these findings, we hypothesized that, along with changes in alpha and beta power, enhanced inhibition after strong visual stimulation would be associated with an acceleration of alpha oscillations.

Another promising but unexplored indicator of the E/I ratio after the cessation of sensory stimulation is the slope of the aperiodic 1/f-like part of the power spectrum. The aperiodic slope is steeper during ‘down states’ such as sleep and anesthesia compared to wakefulness (Colombo et al., 2019; Lendner et al., 2020; Muthukumaraswamy & Liley, 2018), during wakefulness with ‘eyes closed’ compared to ‘eyes open’ (V.O. Manyukhina et al., 2022; F. Zhang et al., 2019) and during awake states characterized by decreased information processing compared to more active states (Ouyang, Hildebrandt, Schmitz, & Herrmann, 2020; Pietrelli, Samaha, & Postle, 2022). The steepness of the aperiodic spectral slope is thought to reflect the neural E/I ratio so that a flatter (less negative) slope is associated with a higher background neuronal firing rates, decoupled from the oscillatory carrier frequency, i.e. with higher ‘noise’ (Gao, Peterson, & Voytek, 2017; Voytek et al., 2015). Given that neuronal firing rate is a hallmark of the E/I ratio in neuronal networks, a steeper (more negative) aperiodic slope in the post-stimulus interval compared to the resting state may be a more direct indicator of decreased excitability than power of periodic alpha-beta oscillations.

To be consistent with the functional inhibition hypothesis, the neurophysiological indices should change in proportion to the strength of the respective sensory input. Indeed, fMRI BOLD and near-infrared spectroscopy (NIRS) oxyhemoglobin undershoots in the visual cortex scale with an intensity of the preceding stimulation (Haigh et al., 2015; Mullinger et al., 2013; Sadaghiani et al., 2009; Stevenson et al., 2011). At the same time, information regarding the influence of the strength of the preceding stimulation on the spatial and spectral features of post-stimulus MEG and EEG is scarce and inconsistent. Stevenson et al analyzed beta MEG power and found greater ‘beta rebound’ after the presentation of drifting vs. static grating in the visual area MT, but not in the primary visual cortex (Stevenson et al., 2011). Mullinger et al (Mullinger et al., 2017), on the other hand, observed a significantly greater increase in EEG alpha power after flickering than after static visual stimulus, but they did not localize the sources of this ‘alpha rebound’ effect.

In the present study, we analyzed MEG recorded during time intervals that followed visual stimulation of varying intensity, as well as during passive visual fixation (‘rest’). We pursued several complementary goals.

*First,* we investigated how various MEG parameters, which had previously been associated with functional inhibition, change after cessation of visual stimulation as compared to the ‘rest’. Apart from the periodic alpha and beta power, we estimated peak alpha frequency (PAF) and the slope of the 1/f part of the power spectrum. There is evidence that the choice of the frequency range for estimating the aperiodic 1/f slope can strongly affect the results (e.g. (V.O. Manyukhina et al., 2022; Maschke, Duclos, Owen, Jerbi, & Blain-Moraes, 2023; Ossandón et al., 2023)). In the majority of the previous studies, the aperiodic slope was estimated either in a wide frequency range, after separating the aperiodic part of the power spectral density (PSD) from the periodic activity, or at high frequencies (>30 Hz), where periodic activity is usually absent. Here we compared these two approaches. First, we used ‘fitting oscillations & one over f’ (FOOOF) algorithm (Donoghue et al., 2020). This method allows separating aperiodic activity from the periodic ‘rhythms’ and is frequently used to estimate the slope in a broad frequency range (e.g. 2-40 Hz). Our second approach was to estimate the aperiodic slope using a linear approximation in the high-frequency range (35-45 Hz) not contaminated by spectral peaks.

*Second*, we sought to find out whether these MEG indices of functional inhibition are proportional to the intensity of the preceding visual stimulation. To modulate the intensity of visual stimulation we varied the drift rate of a high-contrast visual grating that moved toward the center of the visual field. We have previously shown that increasing gratings’ velocity effectively modulates the excitatory state of the visual cortex during the stimulation period, as reflected in changes in high-frequency gamma oscillations (Orekhova et al., 2019; Orekhova et al., 2018) and in the magnitude of pupil constriction (Orekhova et al., 2019).

*Third*, we explored the correlations between the inhibition-sensitive MEG parameters. Although these parameters have been extensively studied individually, the relationship between them is poorly understood. At the same time, the phenomenon of post-stimulus functional inhibition can provide information about this relationship and help elucidate its underlying functional mechanisms.

Our *fourth* aim was to test whether differences in MEG indices of post-stimulus inhibition reflect individual differences in sensory perception. Since visual hypersensitivity is associated with heightened excitability of the visual cortex (Bargary, Furlan, Raynham, Barbur, & Smith, 2015), it may also manifest in an altered post-stimulus inhibition. We considered two possibilities. First, more sensitive individuals may have generally weaker inhibition, which would lead to weaker post-stimulus changes in inhibition-sensitive MEG parameters. Indeed, sensory hypersensitivity in Fragile X Syndrome (FRAX) (Ethridge et al., 2017; Razak, Binder, & Ethell, 2021) and autism (Q. Chen et al., 2020) is thought to reflect a global GABAergic inhibitory deficit. Second, hypersensitive healthy individuals may compensate for increased excitability by down-regulating the excitatory activity of the stimulated sensory areas through inhibitory top-down connections. In line with this hypothesis, it has been shown that subjects with somatic anxiety and elevated sensory sensitivity have increased backward inhibitory effective connectivity in exteroceptive sensory networks (Bouziane, Das, Friston, Caballero-Gaudes, & Ray, 2022). Therefore, if the post-stimulus inhibition is under top-down control, it may be even stronger in hypersensitive individuals. To explore the putative relationship between post-stimulus MEG activity and individual differences in sensory processing, we assessed in our subjects ‘Low Neurological Threshold’ using the Sensory Profile questionnaire (Brown & Dunn, 2002) and investigated whether subjects’ scores on this measure could be predicted by MEG measures of post-stimulus inhibition.

The experimental sample consisted of women who participated in our previous studies devoted to the effect of the menstrual cycle (MC) on visual gamma oscillations (Manyukhina, Orekhova, Prokofyev, Obukhova, & Stroganova, 2022; Manyukhina et al., 2021). Whereas in the previous studies we investigated the direct effect of visual stimulation intensity during stimulus presentation, in the present study we analyzed the time intervals *after* the cessation of these visual stimuli as well as the resting state. The participants were investigated twice, during the follicular and luteal phases of the MC, which provided us with additional information on the role of women’s hormonal status and allowed us to evaluate the between-phase consistency of the metrics used.

## Methods

### 2.1. Participants

Twenty-five healthy females from 18 to 40 years of age (27.9 ± 6.0 years) with normal or corrected vision were recruited for an experiment among participants of the ‘healthy-lifestyle’ internet community or students. Exclusion criteria were the presence of a psychiatric disorder, smoking, irregular MC, and treatment with hormonal therapy. These subjects also participated in our previous study of the effect of MC on brain oscillations (V. O. Manyukhina et al., 2022). Therefore, for each individual, MEG data were collected twice, during the follicular and luteal phases, with an interval between recordings from 7 to 147 days.

The investigations were conducted in accordance with the Declaration of Helsinki and were approved by the Ethical Committee of the Moscow State University of Psychology and Education. All participants provided verbal assent to participate in the study and were informed about their right to withdraw from the study at any time during testing. They also gave written informed consent after the experimental procedures had been fully explained.

### 2.2. Adolescent/Adult Sensory Profile

The participants were asked to fill in the Russian version of the Adolescent/Adult Sensory Profile (A/ASP) (Brown & Dunn, 2002). The A/ASP questionnaire assesses the subject’s sensory processing in daily life according to Brown & Dunn’s four quadrants model: ‘Sensory Sensitivity’, ‘Low Registration’, ‘Sensory Seeking’, and ‘Sensory Avoidance’. In addition, A/ASP allows the assessment of a Low Neurological Threshold, which is calculated as the sum of the Sensory Sensitivity and Sensory Avoidance scales and measures a person’s notice of or annoyance with sensory stimuli of different modalities. Here, to reduce the number of correlations, we used Low Neurological Threshold as an integral measure of subjects’ sensory sensitivity and avoidance and tested how scores on this scale correlate with the MEG indices of post-stimulus inhibition.

### 2.3. MEG experimental design

MEG was recorded under two experimental conditions, at rest and during a visual task (Figure 1A). During rest, subjects fixated their gaze on a small red cross in the center of the screen for five minutes. The visual task is described in detail in our previous studies (Orekhova et al., 2019; Orekhova et al., 2018). In short, the participants were presented with large (18° of visual angle) high-contrast circular grating that remained static (0°/s) or moved with one of three velocities: 1.2°/s (‘slow’), 3.6°/s (‘medium’), 6.0°/s (‘fast’) for 1.2–3 s. Each trial started with a fixation cross which lasted for 1.2 s and was followed by the presentation of a grating of one of the four types in random order. The participants were asked to respond to stimulation changes (motion of initially static grating or cessation of motion of the moving grating) by pressing a button with the right or left index finger (counterbalanced), after which the fixation cross appeared and the next trial began. As the instruction was slightly different for the static and moving stimuli, for the purpose of the present study we only analyzed intervals following moving gratings. The number of omission and commission errors (> 1000 ms and < 150 ms relative to the target event, respectively) was low (mean = 4.1%, std = 2.7%), suggesting that participants attended to the task most of the time. In total, 90 gratings of each type were presented to each individual during three sessions interrupted by short breaks. Short cartoon movies were presented after each 2–5 grating stimuli in order to maintain the subject’s attention and decrease boredom. Figure 1A shows schematic representation of the experimental design.

**Figure 1.**
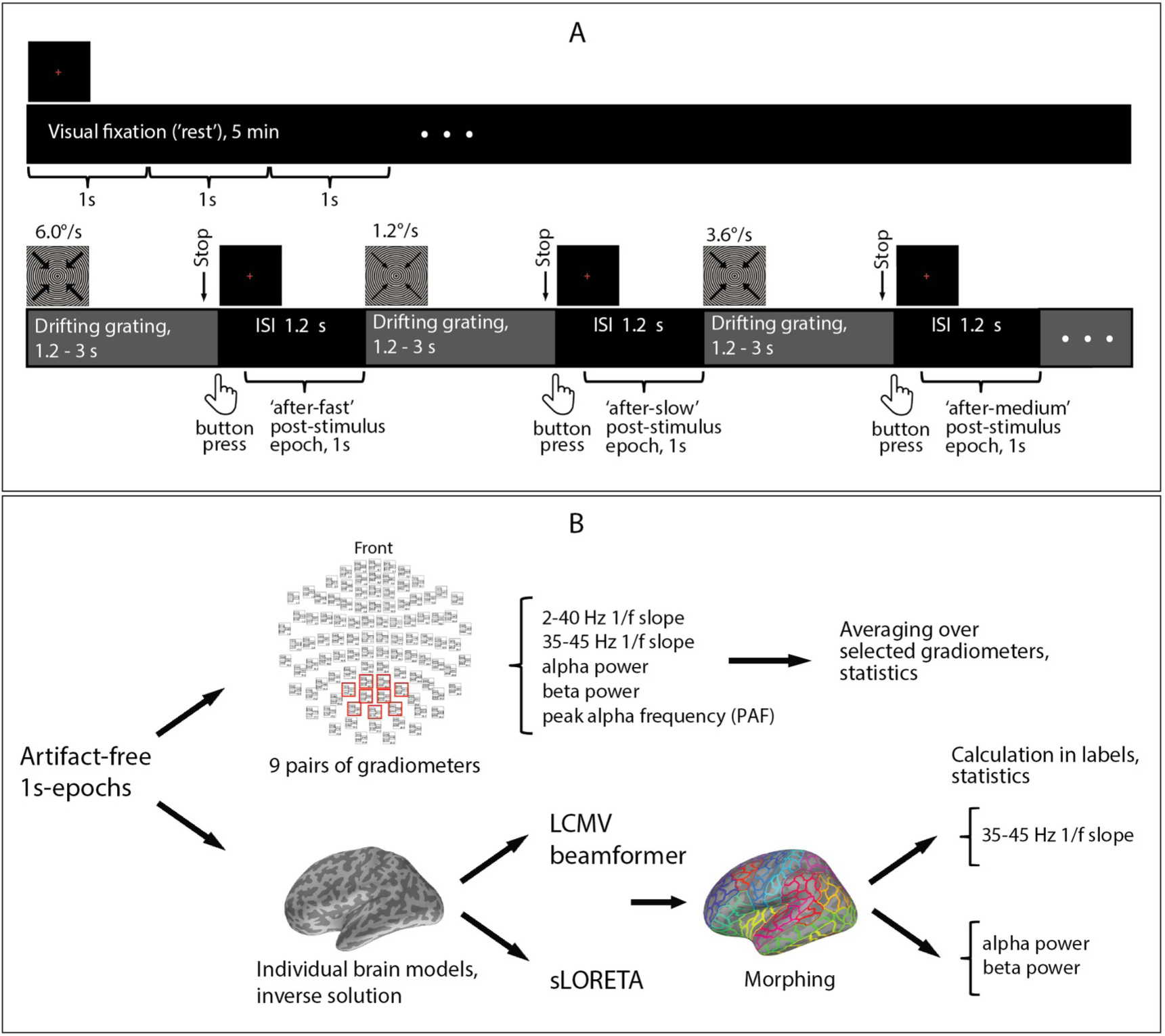
Schematic representation of experimental design and analysis pipeline. (A) MEG was recorded at rest and during a visual task; 1-second epochs after stimulus cessation (marked by brackets) were used for the analysis. (B) Analysis was first performed in the ‘sensor space’, where several MEG parameters were calculated for the selection of posterior gradiometers. Changes in these parameters (except PAF, see Methods) that showed а significant effect of Condition (rest vs. post-stimulus intervals) or of Intensity (‘after-slow’ vs. ‘after-fast’ post-stimulus intervals) were localized in the brain.

### 2.4. MEG data acquisition and preprocessing

MEG data were acquired with an Electa VectorView Neuromag 306-channel MEG detector array (Helsinki, Finland) consisting of 102 magnetometers and 204 planar gradiometers at the Moscow Center for Neurocognitive Research (MEG-center).

Data were registered with 0.03 Hz high-pass and 300 Hz low-pass inbuilt filters and sampled at 1000 Hz. Temporal signal-space separation method (tSSS) (Taulu & Hari, 2009) from MaxFilter software (v.2.2) was applied to the raw MEG signal to reduce external noise and compensate for head movements.

Next preprocessing steps were performed using MNE-python software (v.0.24) (Gramfort et al., 2013). They included data down-sampling at 500 Hz and correction for biological artifacts (heartbeats and vertical eye movements) using Independent Component Analysis (ICA). From 1 to 4 ICs were excluded in each recording (mean±standard deviation (SD): 2.32 ± 0.68 and 2.04 ± 0.34 for the rest and visual task, respectively). The raw data were then filtered at 1-45 Hz to analyze alpha, beta power, and aperiodic 1/f slope estimated using the ‘FOOOF’ algorithm, and at 25-145 Hz to analyze the aperiodic 1/f slope in the 35-45 Hz range (see below). For the rest condition, the raw data were divided into 1s non-overlapping epochs. For the visual task, epochs of 1-s duration were extracted, spanning the period from 0.2 to 1.2 s after cessation of visual stimulation/appearance of the fixation cross (i.e., post-stimulus intervals). Epochs of both types were visually inspected based on unfiltered epoched data, and those contaminated by muscle artifacts or high-amplitude noise were excluded from the analysis. Post-stimulus epochs were further divided according to the type of preceding stimulation (drift rate 1.2°/s, 3.6°/s, or 6.0°/s) into three categories: ‘after-slow’, ‘after-medium’, and ‘after-fast’, respectively. Post-stimulus epochs preceded by cartoon movies were excluded from the analysis. The final number of the clean post-stimulus epochs was 58.76 ± 6.86, 58.04 ± 6.71, and 56.74 ± 6.96 (mean ± SD) for the ‘after-slow’, ‘after-medium’, and ‘after-fast’ conditions, respectively. The number of clean epochs for the rest condition exceeded that for the visual task (all drift velocities combined). To equalize the number of epochs, for each subject, we took the number of clean rest epochs equal to the number of clean post-stimulus epochs from the beginning of the rest condition. For further analysis, we used data from 204 planar gradiometers.

### 2.5. MRI data acquisition and processing

Structural MRI data were acquired in all the participants (General Electric Signa 1.5 Tesla MRI scanner; voxel size 1 mm × 1 mm × 1 mm). T1-weighted anatomical images were processed using the standard ‘recon-all’ pipeline implemented in FreeSurfer software (v.6.0.0) (Fischl et al., 2002) which runs all the default preprocessing steps. For the localization of effects in the brain, the cortical surface was parceled into 448 similar-size labels using anatomical atlas by Khan et al (Khan et al., 2018).

### 2.6. Estimation of aperiodic slopes, alpha and beta power, and peak alpha frequency (PAF) in sensor space

For the sensor space analysis, all MEG data were translated to a standard position (0, 0, 40). Since our visual stimuli induce the strongest neural response in posterior gradiometers (Figure 1B in (Orekhova et al., 2015)), we expected the post-stimulus inhibition to be strongest in the same locations. Therefore, for the analysis at the sensor level we selected 9 pairs of gradiometers, as shown in Figure 1B. We did not include sensors located close to the edges of the MEG helmet in this selection because their signal is frequently contaminated by cranial and neck muscle activity.

We tried two different approaches to estimate the 1/f aperiodic slope from the PSD. First, we estimated the aperiodic slope in the high-frequency range (35-45 Hz), where periodic activity was absent. Second, we estimated it in a broad frequency range (2-40 Hz), after separating periodic and aperiodic components of the power spectrum.

To calculate PSD, we applied Welch’s method (‘psd_welch’ function in MNE-Python software; from 1 to 50 Hz, frequency resolution=0.1 Hz, with zero padding, no overlap) on individual 1-s epochs. The resulting spectra were averaged over epochs and selected gradiometers, separately for the two visits and four conditions: rest and three types of post-stimulus intervals (Figure 1A). The averaged PSD were further used to estimate the aperiodic 1/f slopes in 35-45 Hz and 2-40 Hz ranges, alpha and beta powers and PAF at the sensor level.

To estimate the 35-45 Hz slope, PSD values were interpolated using interpolate.interp1d function from Scipy library in Python v.3.7 in order to have an equal distance between frequency bins on a logarithmic scale. We then fitted the linear function into the logarithm of the PSD vs. the logarithm of frequency using ‘polyfit’ function (Python library NumPy, v.1.21.4), the same as we did in (V.O. Manyukhina et al., 2022).

To estimate the aperiodic slope in the broad frequency range (2-40 Hz), we used the ‘FOOOF’ algorithm (Donoghue et al., 2020). We fitted the model with ‘fixed’ mode (i.e. no ‘knee’) for the ‘aperiodic_mode’ parameter, all other parameters were set to default (peak_width_limits=(0.5,12), max_n_peaks=inf, min_peak_height=0, peak_threshold=2). To obtain a slope coefficient comparable to the one estimated for the 35-45 Hz, the aperiodic exponent output of the FOOOF was multiplied by -1. We also attempted to fit the model with a ‘knee’ mode (aperiodic_mode=’knee’; all other parameters were the same). The results of this fit are provided in Supplementary Results.

FOOOF was also used to estimate the periodic component of the power spectrum:

> Periodic = Original spectrum - Spectrum without periodic peaks,

where ‘Spectrum without periodic peaks’ was calculated using model._spectrum_peak_rm FOOOF function. This periodic component was then used to calculate mean alpha and beta power as the average power in frequency bands 8-14.3 Hz and 15-25 Hz, respectively. To distinguish alpha frequency changes from changes in alpha-beta power, we estimated the frequency of the highest peak in the alpha band (i.e., peak alpha frequency, PAF) rather than the weighted alpha frequency. The peak in PSD was identified using ‘signal.find_peaks’ function from Scipy library v.1.7.3). As the alpha peak was present in all the conditions only in 22 subjects, PAF was calculated in 22 of 25 women.

### 2.6. Source space analysis

For each visit session and for each condition, the data were corrected to the subject’s ‘optimal’ initial head position (see (Orekhova et al., 2023) for details). Individual structural MRIs were co-registered with the subject’s MEG recordings using ‘mne_analyse’ tool from MNE-C software (https://mne.tools/stable/install/mne_c.html). Further steps of source-level analysis were performed using MNE-python software (v.0.24) (Gramfort et al., 2013). For each subject, a single-layer boundary element model and surface-based source space with 4098 vertices in each hemisphere were created, and a forward solution was estimated.

To estimate the FOOOF-based parameters (aperiodic slope in 2-40 Hz range, alpha and beta power), the raw data were filtered at 1-45 Hz, and standardized low resolution brain electromagnetic tomography (sLORETA) inverse solution was computed. For the analysis of 35-45 Hz 1/f slope, the raw data were filtered at 25-145 Hz. In the latter case, to solve the inverse problem we used the linearly constrained minimum variance (LCMV) beamformer. The choice in favor of different inverse solution methods was made based on their different advantages and disadvantages. LCMV beamformer, together with the broad-band filtration in the high-frequency range, is preferable for analysis of the 35-45 Hz 1/f slope because it enables better filtering of noise and myogenic artifacts than sLORETA (V.O. Manyukhina et al., 2022). sLORETA, on the other hand, does not have depth bias in the source localization and therefore provides more realistic cortical localization of spectral power than LCMV beamformer (Tait, Özkan, Szul, & Zhang, 2021).

Filtered raw data were epoched, and bad epochs (see Section 2.4) were discarded. Different spatial filters were calculated for the two types of comparisons. To compare post-stimulus intervals that followed different types of stimulation (‘after-slow’, ‘after-medium’, ‘after-fast’), the common spatial filter was calculated for these three conditions. For the comparison between rest and post-stimulus intervals pooled across stimulus types, a common spatial filter was calculated for all of these data. For the source reconstruction with the LCMV beamformer, the noise covariance parameter in the filter estimation function was set to ‘None’, which was possible because the analysis was performed for one type of sensors (gradiometers). The unit-gain LCMV beamformer filter was created with the orientation that maximizes power, regularization 0.1, and the ‘reduce_rank’ parameter set to ‘True’. For sLORETA, noise covariance was estimated from the empty room MEG data, which were preprocessed in the same way as the raw data. ‘Loose’ and ‘depth’ parameters were set to 0.4 and 0.8, respectively. The LCMV beamformer filter or the sLORETA inverse operator was applied to each epoch individually.

MEG parameters in the source space were estimated in the same way as those in the sensor space, but in this case, PSD was calculated at the level of vertex sources and averaged within each of 448 cortical labels chosen according to an anatomical parcellation (Khan et al., 2018). To estimate the 35-45 Hz aperiodic slope, the PSD at each individual vertex was normalized to its maximal value before averaging. This was done to ensure that all vertices contributed equally to the average. We did not analyze PAF in the source space, because many participants lacked distinct alpha peaks beyond parieto-occipital cortical areas.

### 2.8. Statistical analysis

To estimate the reliability of the inhibition-sensitive MEG variables between phases of the MC we calculated intraclass correlation coefficients (ICCs). ICCs were estimated for a fixed set of ‘raters’ (follicular and luteal MC phases) (Koo & Li, 2017).

For the sensor-level analysis, all variables were checked for deviation from a normal distribution using Shapiro-Wilk’s W test. Alpha and beta spectral power values were preliminary log10-transformed to normalize the distributions. The resulting distributions did not differ significantly from the Gaussian. Levene’s test was used to test for homogeneity of variance. As none of the analysis of variance (ANOVA) assumptions was violated, we then used repeated-measures ANOVAs (rmANOVAs) to test for the effects of Condition *or* Intensity and Phase (follicular, luteal), as well as for the interaction effects. Factor Condition had 2 levels (rest and post-stimulus period pooled across all trials), while the factor Intensity had 3 levels (‘after-slow’, ‘after-medium’, and ‘after-fast’). When appropriate, the Greenhouse-Geisser correction was applied to correct for violation of the sphericity assumption.

For the source-space analysis, in each of the 448 cortical labels, we used Student’s t-test for related samples to test for the difference between rest and post-stimulus condition or between post-stimulus intervals following cessation of the most and least intensive stimulation (i.e. ‘after-fast’ vs ‘after-slow’ intervals). To control for multiple comparisons, we applied Benjamini-Hochberg false discovery rate (FDR) correction with a threshold of 0.05 to the p-values.

To test for a relationship between condition- or intensity-related changes in MEG parameters and Low Neurological Threshold we estimated Pearson correlation coefficients. In this case, changes in spectral power between conditions or intensities were quantified in %, as 2*(A-B)/(A+B)*100%, while changes in PAF and aperiodic slopes were calculated as the absolute differences: A - B.

## 3. Results

To investigate the sensitivity of MEG parameters to post-stimulus enhancement of inhibition, we followed two lines of analysis. *First*, we compared time intervals following cessation of visual stimulation (all three types of post-stimulus epochs combined) with the resting state, when subjects passively fixated their gaze on a central cross for a long period of time (Figure 1A). We expected to find concordant changes in aperiodic slope, alpha-beta power, and PAF that would indicate enhancement of inhibition during the post-stimulus period compared to rest. *Second*, to find out whether the inhibition-sensitive MEG parameters change in proportion to the intensity of the preceding stimulation, we compared post-stimulus intervals that followed visual stimulation of increasing intensity: gratings drifting at ‘slow’, ‘medium’, and ‘fast’ rates.

We started with an analysis in the ‘sensor space’. In this case, each of the parameters was averaged over a selection of 9 pairs of posterior gradiometers (Figure 1B) and subjected to rmANOVA to test for the effects of stimulation condition or intensity. In addition, for each parameter, we estimated the ICC between the two phases of the MC. We then analyzed the cortical localization of significant condition- and intensity-related differences found at the sensor level.

### 3.1. Sensor space analysis

#### Intraclass correlations (ICCs) of the MEG parameters

To find out whether the investigated parameters demonstrated rank-order stability across the two phases of the MC, we calculated ICCs between the two visits (during follicular and luteal phases of the MC). Nearly all ICCs were good (>0.75) (Koo & Li, 2017) suggesting high stability (Table 1). It should be noted that a low ICC does not necessarily indicate low stability, as it may reflect a systematic but individually specific effect of MC.

**Table 1.**
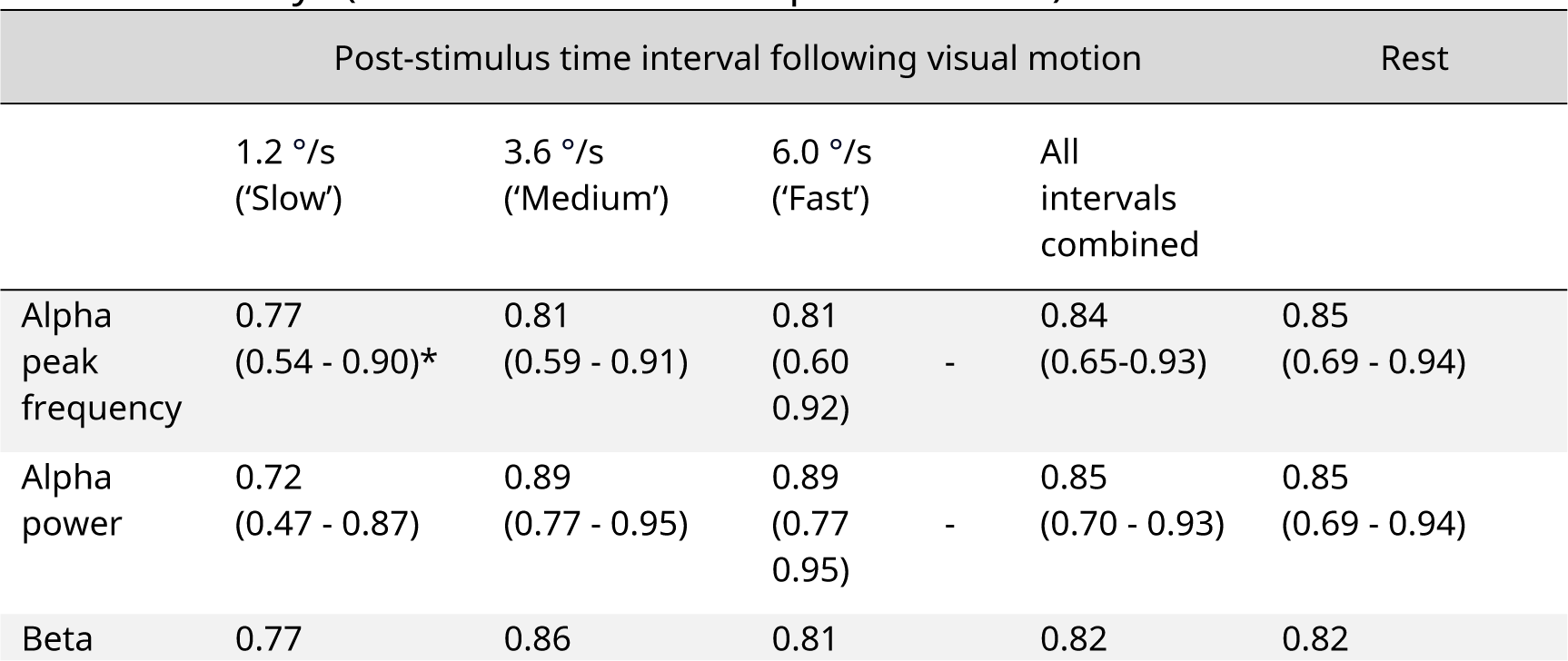
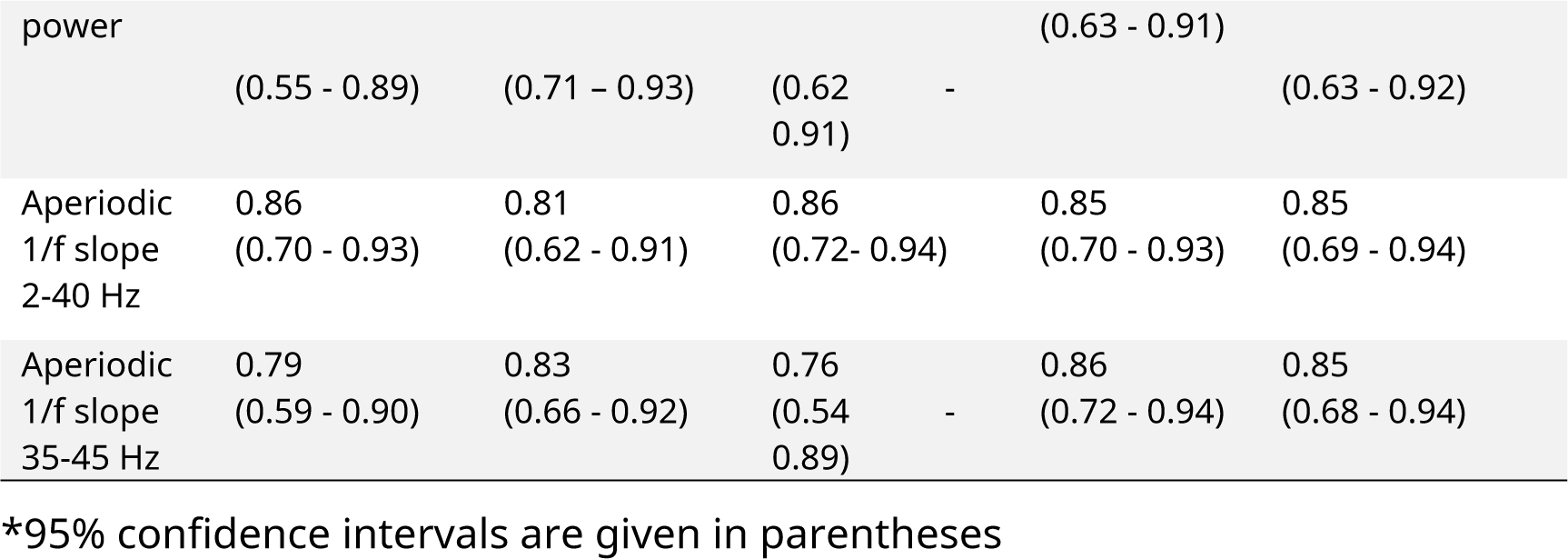
ICCs of the MEG parameters between the recording sessions performed on different days (in follicular and luteal phases of MC).

#### Comparison of aperiodic slopes estimated by two different methods

The aperiodic slope estimated by FOOOF in the 2-40 Hz range was significantly flatter than that estimated by linear approximation of the log-transformed spectrum in the high-frequency (35-45 Hz) range (T-test for dependent samples: for all conditions and visits T(24)>5.6, p<0.00001). Supplementary figure S1 shows slope regression lines calculated using the two methods. There were no significant correlations between 2-40 Hz and 35-45 Hz aperiodic slope coefficients (all correlations calculated for different MC phases and conditions were *negative*; p’s > 0.05).

Next, we tested whether the aperiodic slopes estimated with two different methods showed the previously described relationship with the periodic activity (Muthukumaraswamy & Liley, 2018; Podvalny et al., 2015). Since both alpha and beta oscillations are associated with a reduction of neuronal activity (Iemi et al., 2022), here we tested for correlation of aperiodic slopes with both alpha and beta periodic power. The aperiodic 1/f slope estimated in the 35-45 Hz frequency range indeed negatively correlated with alpha and beta periodic power during both rest and post-stimulus period (Table 2). No significant correlations were observed for the slope estimated in the 2-40 Hz range using FOOOF (Table 2). PAF did not correlate with either of the aperiodic slopes (Table 2).

**Table 2.** Pearson correlations of aperiodic spectral slope (estimated by two different methods) with periodic alpha power, periodic beta power, and peak alpha frequency (PAF).

|  | Rest |  |  | Post-stimulus time interval |  |  |
| --- | --- | --- | --- | --- | --- | --- |
|  | Periodic alpha power (N=25) | Periodic beta power (N=25) | PAF (N=22) | Periodic alpha power (N=25) | Periodic beta power (N=25) | PAF (N=22) |
| Aperiodic 1/f slope 35-45 Hz |  |  |  |  |  |  |
| *Follicular phase* | **-0.66\*\*** | **-0.60\*\*** | 0.29 | **-0.68\*\*** | **-0.67\*\*** | 0.30 |
| *Luteal phase* | **-0.62\*\*** | **-0.61\*\*** | 0.29 | **-0.66\*\*** | **-0.69\*\*** | 0.21 |
| Aperiodic 1/f slope 2-40 Hz |  |  |  |  |  |  |
| *Follicular phase* | 0.23 | 0.26 | 0.21 | 0.06 | 0.12 | 0.20 |
| *Luteal phase* | 0.28 | 0.35 | 0.06 | 0.25 | 0.35 | -0.04 |

N – number of subjects; \*\* p<0.004, FDR corrected

A recent MEG study showed that the MEG aperiodic activity is better approximated by two different linear functions with a ‘knee’ at approximately 15 Hz (Ibarra Chaoul & Siegel, 2021) Therefore we also fitted FOOOF using a ‘knee mode’. As a result of this analysis, the knee varied between 2 and 26 Hz for different subjects and conditions, and the post-knee aperiodic slope coefficient was highly variable (Supplementary Results, figures S2, S4). In some cases, the knee frequency markedly differed between different recording sessions for the same subject and condition (Supplementary results, figure S3). Considering these cofounds, we report here only the results obtained using FOOOF with ‘no knee’ mode.

#### Comparison between rest and post-stimulus period

Figure 2A shows the power spectra for the two experimental conditions - rest and period after cessation of visual stimulation averaged for all participants over 9 pairs of posterior gradiometers and two visits across all participants.

**Figure 2.**
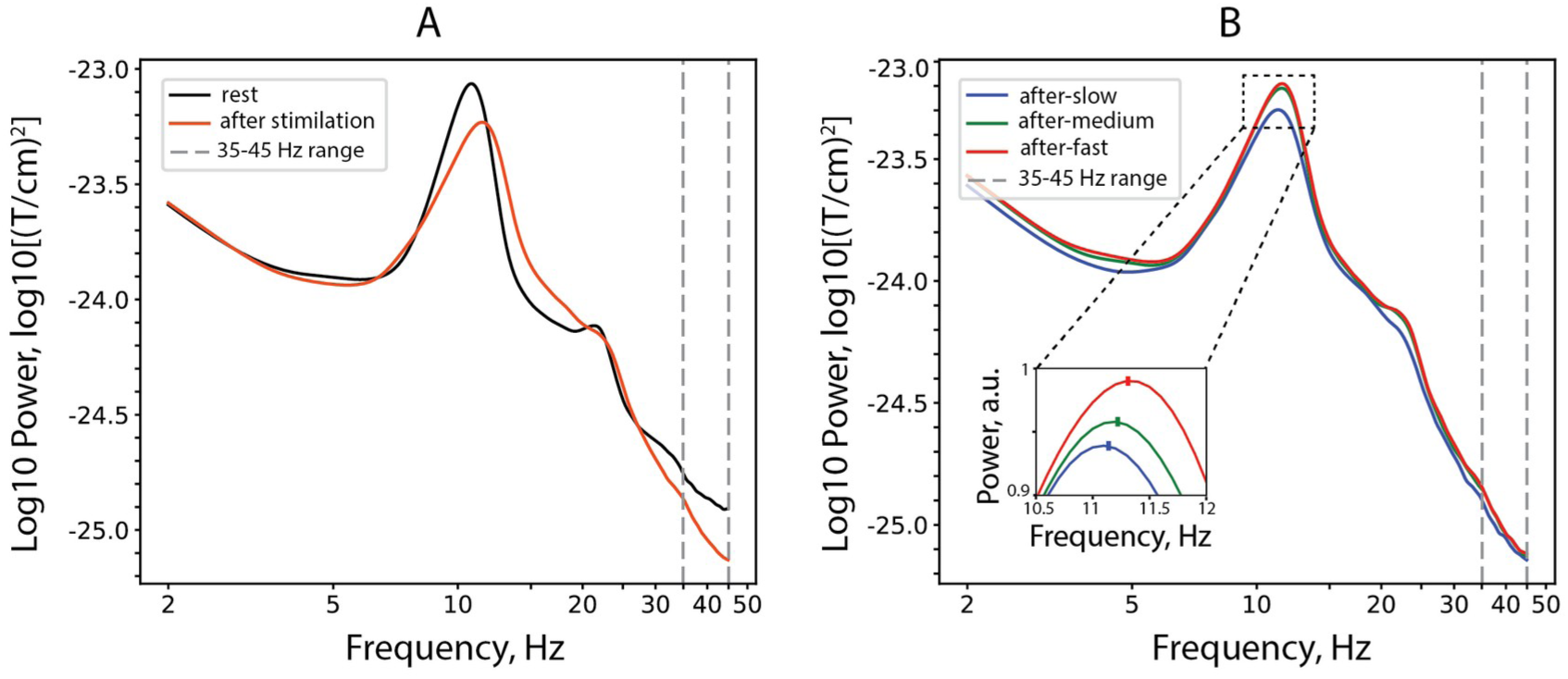
Grand-average log-transformed power spectra for the rest and post-stimulus conditions. (A) Rest and combined post-stimulus intervals. (B) Intervals after cessation of stimulation with slow-, medium-, and fast-drifting gratings. The insert that zooms on the alpha peak shows relative power (i.e., each frequency bin is normalized by the sum over the 2-40 Hz range and multiplied by 100). Only subjects with a distinct alpha peak in all conditions are included (N=22). Note that post-stimulus PAF (marked with a dash in the insert) increases proportionally to the drift rate of the preceding grating.

Comparison between MEG parameters estimated during rest and post-stimulus intervals are shown in Figure 3A. The rmANOVAs with Condition (rest, post-stimulus) and Phase (follicular, luteal) factors showed that the 35-45 Hz slope was steeper during the post-stimulus period (F(1,24)=13.8, p=0.001; η_p_^2^ = 0.35), while no such effect was found for the 2-40 Hz 1/f slope (F(1,24)=0.05, p=0.83). We assumed that if the condition-related differences in the 35-45 Hz aperiodic slope in our study are related to muscle artifacts that are typically evident in MEG at frequencies above 20 Hz (Muthukumaraswamy, 2013), then they should be more pronounced at edge gradiometers closer to the neck muscles (Supplementary Results, figure S6). However, additional analysis (see Supplementary Results) revealed no significant effect of Condition (or Intensity) for edge gradiometers, suggesting that the observed effect is unlikely to be explained by between-condition differences in muscle artifacts.

**Figure 3.**
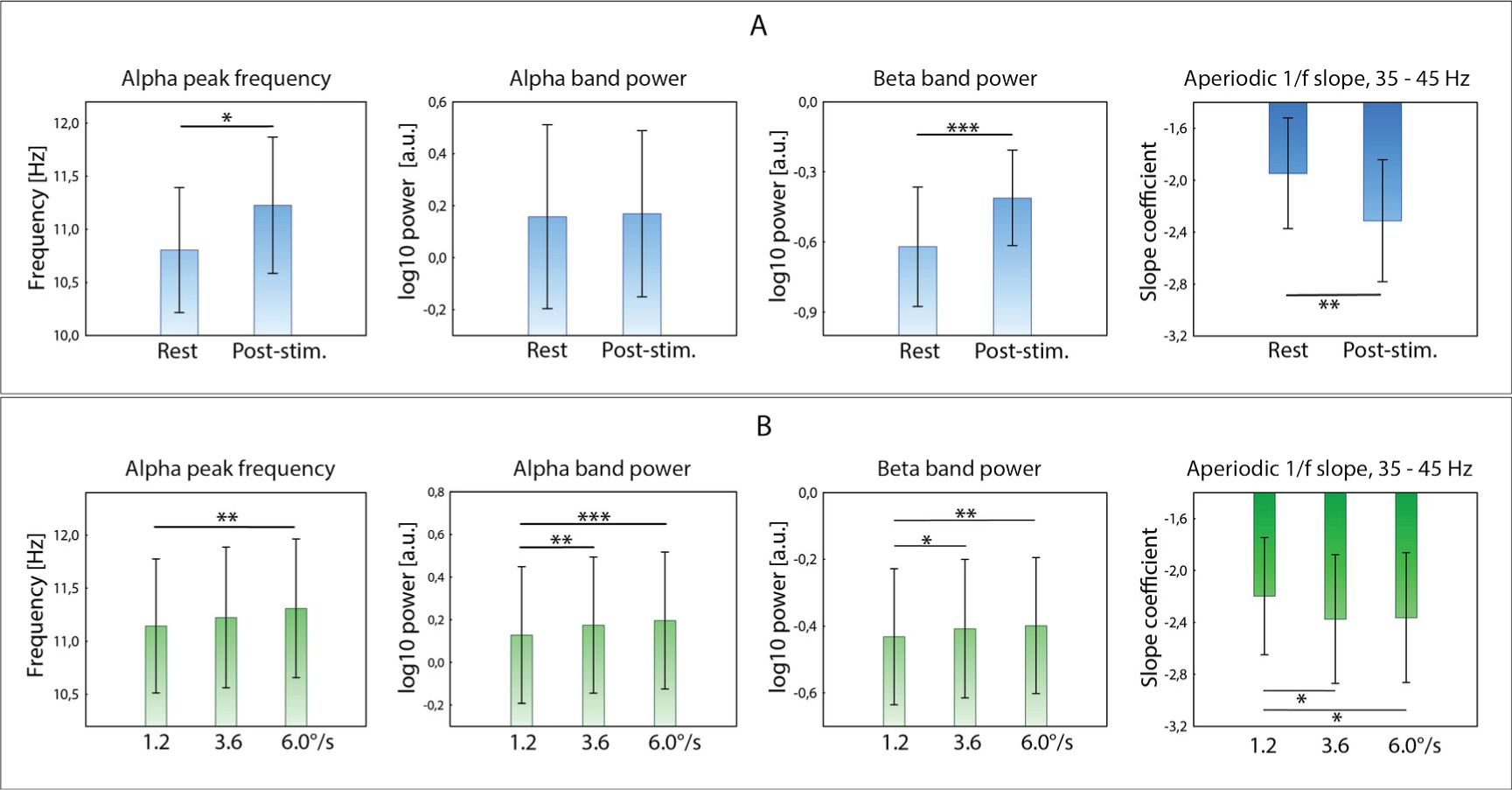
Effects of Condition and stimulation Intensity on MEG indices of functional inhibition. MEG parameters were averaged over 9 pairs of posterior gradiometers. (A) Comparison of conditions: rest vs. post-stimulus interval. (B) Comparisons of post-stimulus intervals as a function of preceding stimulation intensity (i.e., grating’s drift rate).

A spectral alpha peak in all conditions was present in 22 of 25 participants. In these participants, PAF increased during the post-stimulus period relative to rest (Mean_rest_ = 10.8, SD = 0.28; Mean_post_ = 11.2, SD = 0.31; F(1,21)=8.0, p=0.01; η_p_^2^ = 0.27).

Periodic beta power strongly increased during the post-stimulus period compared to rest (F(1,24)=27.9, p=0.00002; η_p_^2^ = 0.54), while no condition-related differences were observed for periodic alpha power (F(1,24)=0.03, p=0.87). A lack of alpha power reactivity to a putative increase of inhibition in the ISI might be explained, at least in part, by the counteracting effect of increased post-stimulus inhibition and increased tonic arousal on alpha power. In subjects with a low level of tonic arousal in a calm resting state condition, a visual task requiring attention can elicit an alerting effect. In such subjects, the level of arousal during post-stimulus task intervals may be higher than during the resting period, resulting in a decrease in alpha power. Consistent with this hypothesis, subjects with higher than average resting alpha power levels (and likely low arousal level in this condition) also showed a significant decrease in alpha power during post-stimulus period (Wilcoxon signed-rank test: Follicular phase, N=7, Z=2.0, p=0.04; Luteal phase: N=9, Z=2.1, p=0.04), while no such effect was observed in subjects with below-average resting-state alpha power (both p’s>0.6).

The effect of the MC Phase was significant only for PAF: alpha frequency was higher in the luteal than in the follicular phase (follicular Mean = 10.9, SD = 0.27; luteal Mean = 11.2, SD = 0.29; F(1,21)=12.9, p=0.002; η_p_^2^ = 0.38).

#### Effect of the intensity of the preceding stimulation

Figure 2B shows the group mean power spectra (averaged over 9 pairs of posterior gradiometers and two visits) for time intervals after the disappearance of visual gratings, separately for three types of stimuli. Figure 3B shows the comparison between time intervals that followed the cessation of gratings drifting at different rates. The rmANOVAs with factors Intensity (‘after-slow’, ‘after-medium’, ‘after-fast’) and Phase (follicular, luteal) showed a significant main effect of Intensity on post-stimulus alpha power (F(2,48) =15.2, p=0.00001; η_p_ ^2^ = 0.39), beta power (F(2,48)=6.9, p=0.002; η_p_ ^2^ = 0.22), PAF (F(2,42)=4.5, p=0.02; η_p_ ^2^ = 0.18), and 35-45 Hz 1/f aperiodic slope (F(2,48)=4.1, p=0.02; η_p_ ^2^ = 0.15). Alpha power, beta power, and PAF increased, while the 35-45 Hz 1/f slope became steeper as the intensity of the preceding stimulation increased. No intensity-related changes in the 2-40 Hz slope were detected (F(2,48)=0.51, p=0.6).

Again, the effect of Phase was significant only for the PAF (follicular Mean = 11.10, SD = 0.40; luteal Mean = 11.4, SD = 0.39; F(1,21)=8.5, p=0.008; η_p_ ^2^ = 0.29). There were no significant Condition x Phase interactions.

#### Correlations between condition-related changes in MEG indices of functional inhibition

To assess whether changes in the MEG parameters sensitive to inhibition reflect similar underlying mechanisms, we calculated correlations between their changes (Tables 3A and 3B). Increasing steepness (negativity) of 35-45 Hz 1/f slope from ‘after-slow’ to ‘after-fast’ post-stimulus interval correlated (or tended to correlate) with an increase in alpha-beta power. Changes in PAF did not correlate with changes in other parameters even as a tendency.

**Table 3A.** Correlation of changes in inhibition-sensitive MEG parameters between rest and post-stimulus period.

|  | Peak alpha frequency<br>Post-stimulus - Rest | Periodic alpha power<br>Post-stimulus/Rest | Periodic beta power<br>Post-stimulus/Rest |
| --- | --- | --- | --- |
| Periodic alpha power<br>Post-stimulus/Rest | -0.13<br>(N=22) |  |  |
| Periodic beta power<br>Post-stimulus/Rest | -0.11<br>(N=22) | **0.56\***<br>(N=25) |  |
| 1/f slope 35-45 Hz<br>Post-stimulus - Rest | -0.10<br>(N=22) | -0.36<br>(N=25) | -0.41<br>(N=25) |

\* $p < 0.05$ , FDR corrected. Here and hereafter, the parameters were averaged over 2 visits.

**Table 3B.** Correlation of changes in inhibition-sensitive MEG parameters between ‘after-slow’ and ‘after-fast’ conditions.

|  | Peak alpha frequency<br>'after-fast' - 'after-slow' | Periodic alpha power<br>'after-fast'/'after-slow' | Periodic beta power<br>'after-fast'/'after-slow' |
| --- | --- | --- | --- |
| Periodic alpha power<br>'after-fast'/'after-slow' | 0.06<br>(N=22) |  |  |
| Periodic beta power<br>'after-fast'/'after-slow' | -0.34<br>(N=22) | **0.76\*\*\***<br>(N=25) |  |
| 1/f slope 35-45 Hz<br>'after-fast' - 'after-slow' | 0.07<br>(N=22) | -0.40#<br>(N=25) | **-0.48\***<br>(N=25) |

To sum up, the sensor-level analysis suggested that the MEG parameters putatively sensitive to functional inhibition changed from rest to post-stimulus condition and scaled with an intensity of preceding stimulation. The increase of post-stimulus inhibition as compared to rest was manifested by an increase in beta power, alpha frequency, and the steepening of the 1/f aperiodic slope estimated in the 35-45 Hz frequency range. The scaling of post-stimulus inhibition with the increase of the preceding stimulation intensity was reflected in concordant changes in the four investigated parameters: increase of alpha and beta power, increase of PAF, and steepening of the 35-45 Hz aperiodic slope in the post-stimulus intervals following the ‘fast’ vs. ‘slow’ drift rate of moving visual gratings. The aperiodic slope estimated in the 2-40 Hz range with FOOOF did not correlate with that estimated in the 35-45 Hz frequency range or with alpha-beta power and did not vary from rest to post-stimulus condition or with the intensity of the preceding visual input.

### 3.2. Source space analysis

To find cortical sources of the significant effects observed at the sensor level for alpha and beta power, as well as for the 35-45 Hz aperiodic slope, we performed a source localization analysis. Although PAF also demonstrated significant condition- and intensity-related changes, we did not perform this analysis for PAF, because many participants lacked distinct alpha peaks outside the posterior cortical areas.

Figure 4 shows the cortical distribution of alpha and beta periodic power, as well as that of 1/f aperiodic slope at rest and in the post-stimulus period (all drift rates of visual gratings combined). Figure 5 shows cortical regions where differences in spectral power and aperiodic slope between rest and post-stimulus conditions were significant.

**Figure 4.**
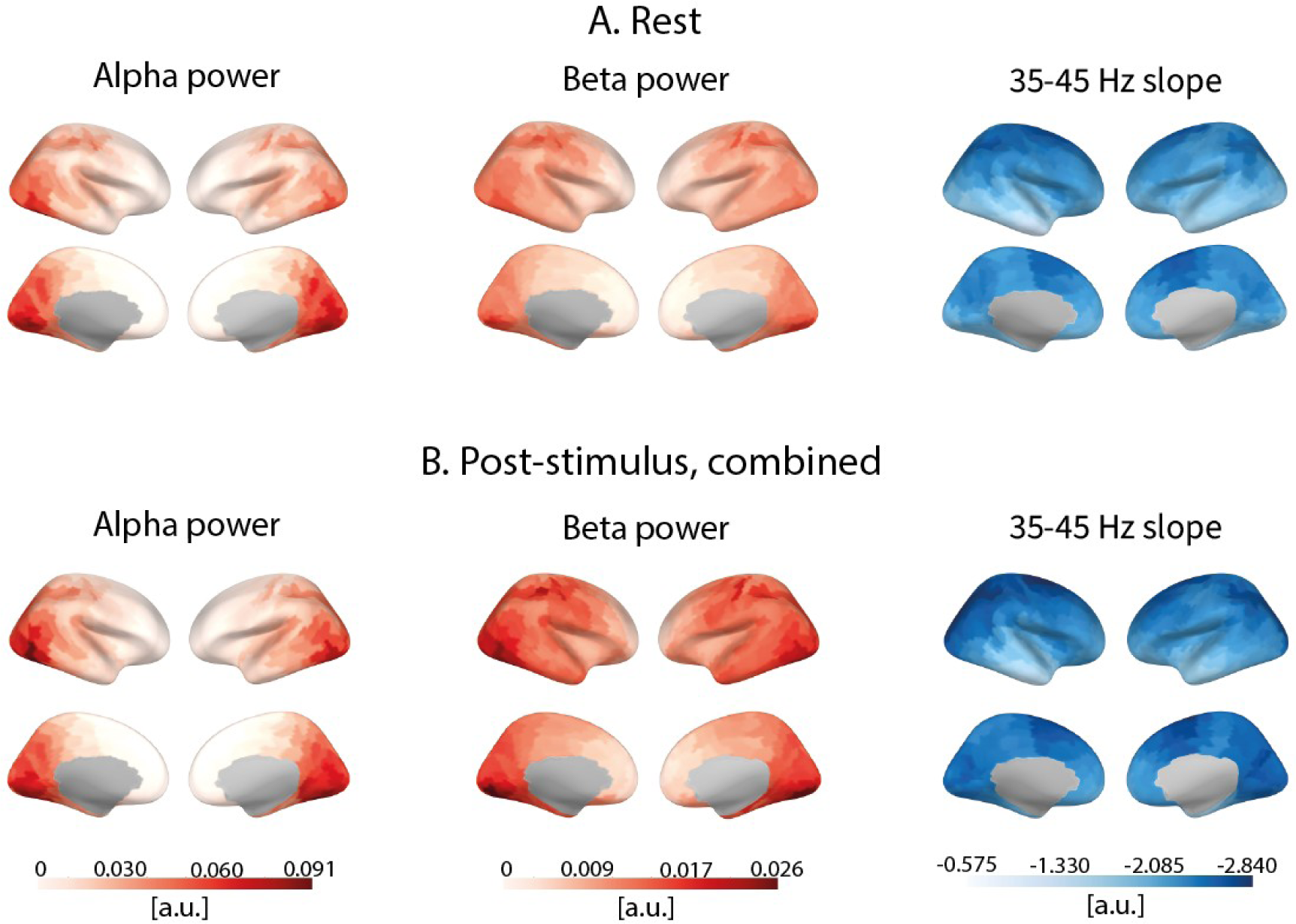
Cortical distribution of periodic alpha power, periodic beta power, and 1/f aperiodic slope estimated in 35-45 Hz range during (A) rest and (B) post-stimulus intervals combined over the three types of the preceding visual stimuli.

**Figure 5.**
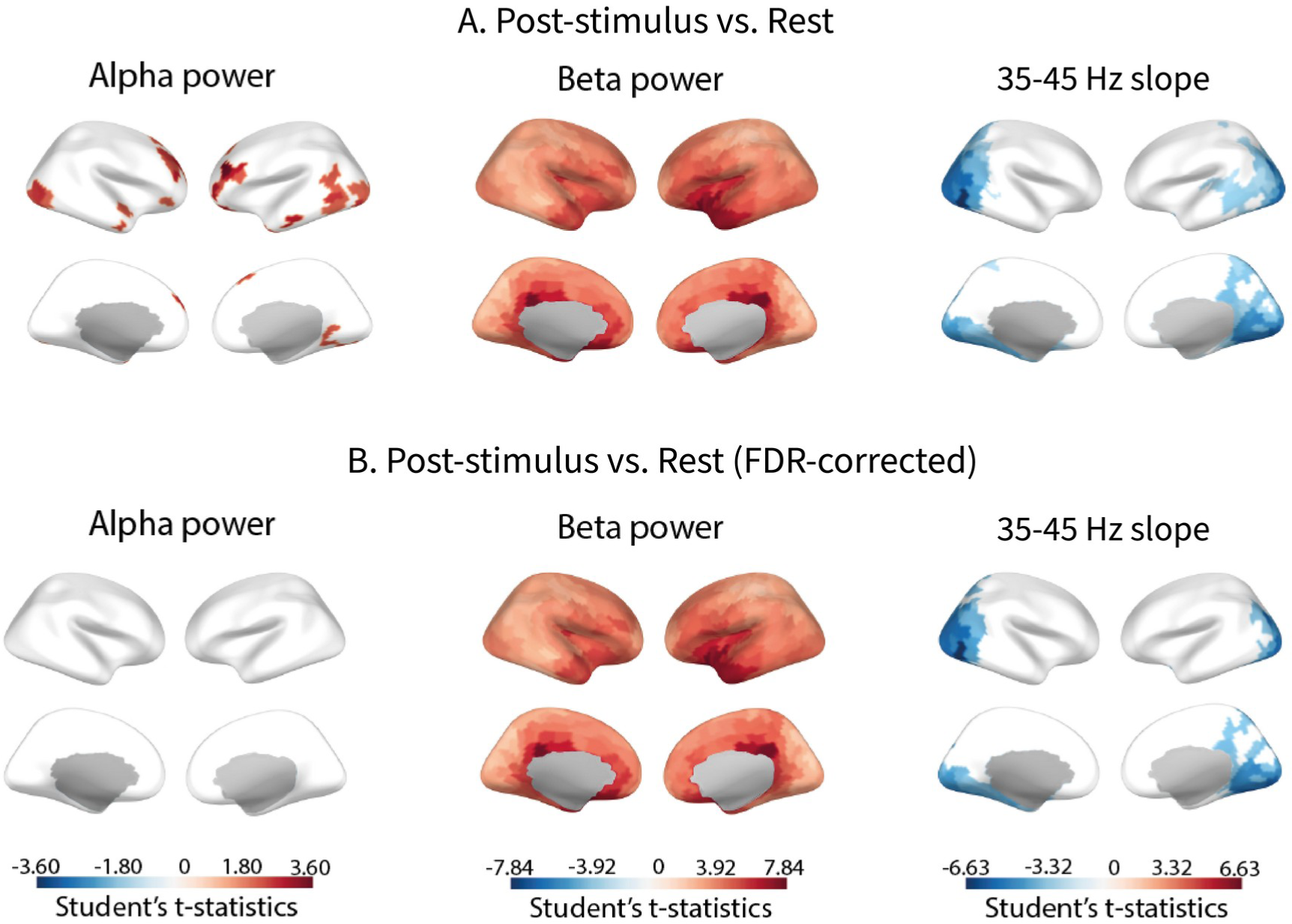
Statistical maps of difference in inhibition-sensitive MEG parameters between rest and post-stimulus intervals, pooled for ‘after-slow’, ‘after-medium’, and ‘after-fast’ trials. On the color-coded t-values scale, significant increases and decreases in parameter values from rest to post-stimulus condition are indicated in red and blue, respectively. (A) Differences significant (p<0.05) without correction for multiple comparisons. (B) Differences significant (p<0.05) after FDR correction.

The decrease in the aperiodic spectral slope from the rest to post-stimulus interval was observed in the posterior areas, including primary and secondary visual cortex. Unlike that for the slope, differences in the beta power between rest and post-stimulus conditions were widespread and most significant in the cingulate cortex, orbitofrontal cortex, and insula.

Figure 6 illustrates differences in the MEG parameters between ‘after-fast’ vs. ‘after-slow’ conditions. The period after the fast-moving stimulus (more intense stimulation) compared to the period after the slow-moving stimulus (less intense stimulation) was characterized by greater alpha and beta power in visual cortical areas. For alpha power, the difference was also observed in the right precuneus, posterior cingulum, and retrosplenial cortex. The 35-45 Hz aperiodic slope was more negative after more intense stimulation in the right supramarginal area and bilaterally in the visual cortex. Although the stimulation intensity-related changes of the 35-45 Hz aperiodic slope did not survive correction for multiple comparisons at the source level, given the significant results at the sensor-level (Figure 3B), we considered these changes to be reliable.

**Figure 6.**
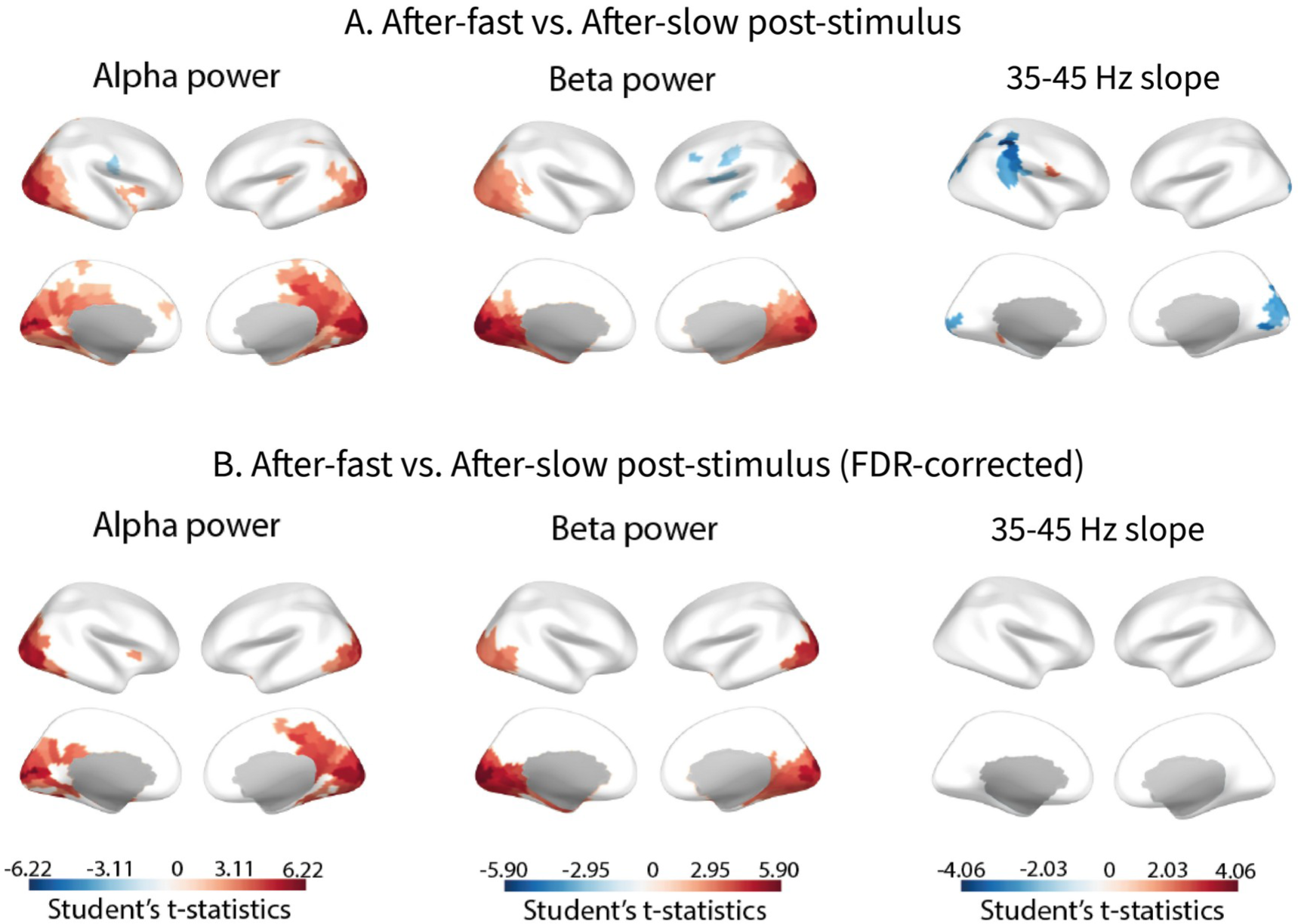
Statistical maps of the differences between ‘after-slow’ (drift rate 1.2 °/s) and ‘after-fast’ (drift rate 6.0 °/s) post-stimulus intervals. On the color-coded t-values scale, significantly higher and lower parameter values in the ‘after-fast’ compared to ‘after-slow’ intervals are indicated in red and blue, respectively. (A) Differences significant (p<0.05) without correction for multiple comparisons. (B) Differences significant (p<0.05) after FDR correction.

To sum up, results at the source level showed that in comparison with rest, the period following cessation of visual stimulation was characterized by a broad increase in beta power and steepening of the aperiodic slope in the visual cortical areas. Increases in alpha and beta power and steepening of the 35-45 Hz aperiodic slope after more intense compared with less intense visual stimulation were observed more locally in the visual cortex and nearby areas.

### 3.3 The link between post-stimulus inhibition and Low Neurological Threshold

We expected that inter-individual differences in post-stimulus inhibition might be related to individual variability in subjective sensory sensitivity and avoidance. To test this prediction, we calculated correlations between subjects’ scores on the Low Neurological Threshold and condition- or intensity-related changes in inhibition-sensitive parameters (PAF, alpha power, beta power, 35-45 Hz aperiodic slope) (Table 4). Only those parameters, which demonstrated significant condition- or intensity-related differences at the sensor level were used for this analysis (Figures 3A,B).

**Table 4.** Pearson correlations between Low Neurological Threshold scores and condition- or intensity-related changes in MEG parameters sensitive to post-stimulus inhibition.^#^.

|  | Post-stimulus intervals vs. rest |  |  | 'After-fast' vs. 'after-slow' |  |  |  |
| --- | --- | --- | --- | --- | --- | --- | --- |
|  | PAF, difference | Beta power, % change | 35-45 Hz slope, difference | PAF, difference | Alpha power, % change | Beta power, % change | 35-45 Hz slope, difference |
| Low Neurological Threshold | R=0.44 | R=0.09 | R=-0.31 | R=-0.24 | **R=0.57\*** | R=0.52 | R=-0.19 |

<sup>#</sup> All the parameters were estimated at the sensor level. Changes in PAF and aperiodic slope were calculated as the absolute difference between conditions, while changes in periodic power were calculated as the relative difference in % (see Methods for details). Three subjects lacking alpha peaks in one or more conditions and one participant lacking data on Low Neurological Threshold were excluded from this analysis leaving a total of 21 subjects. \*p= 0.042, FDR-corrected.

Higher scores on the Low Neurological Threshold (i.e., greater sensitivity to sensory stimulation and avoidance of it in daily life) correlated with greater intensity-related changes between ‘after-slow’ and ‘after-fast’ conditions in alpha (p=0.042, FDR-corrected) and - as a tendency - beta (p=0.06, FDR-corrected) power (Figures 7A,B).

**Figure 7.**
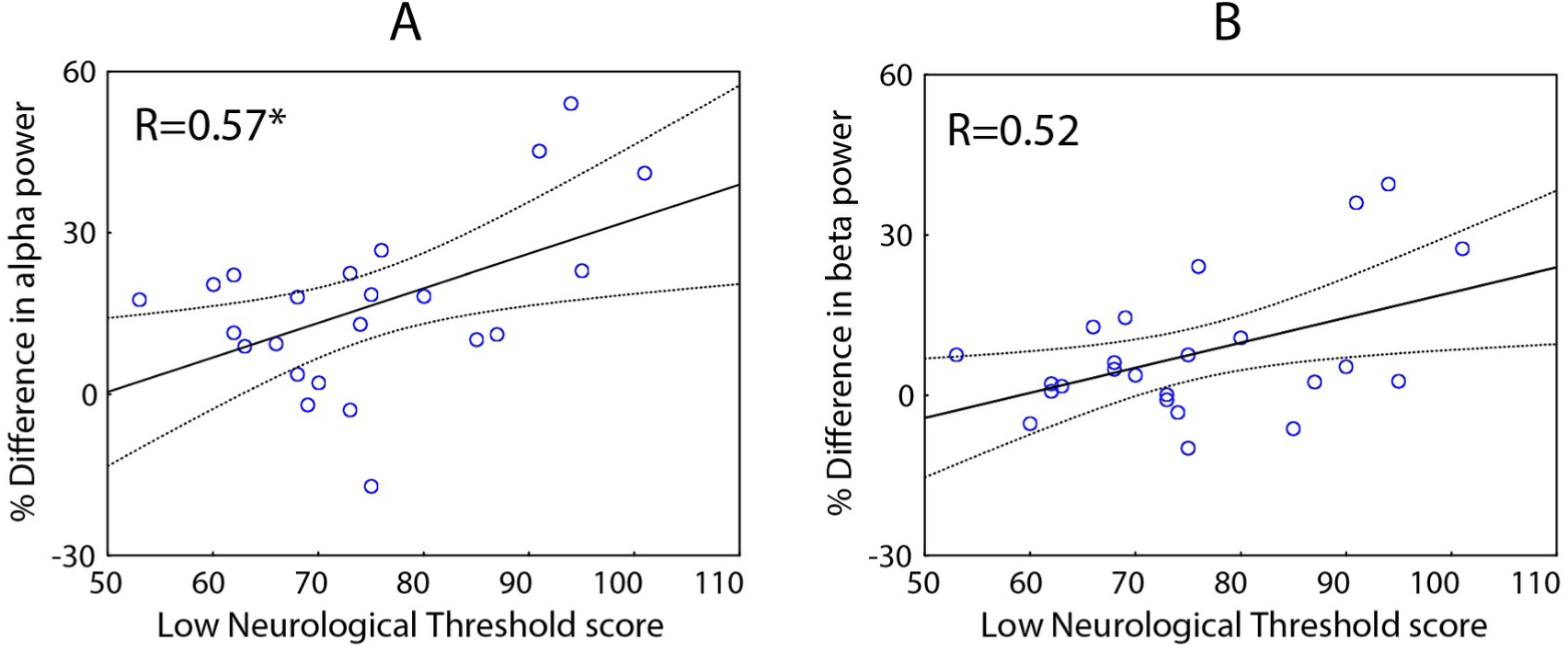
The relationship between intensity-related increase in periodic alpha (A) and beta (B) power from ‘after-slow’ to ‘after-fast’ condition and Low Neurological Threshold (individual sensory sensitivity and avoidance) scores. The relative difference between ‘after-fast’ and ‘after-slow’ post-stimulus intervals is calculated for the sensor space data, as 2*(after-fast – after-slow)/(after-fast + after-slow)*100%.

## 4. Discussion

The neuronal studies suggest that cessation of sensory stimulation is followed by an increase in functional inhibition, which exceeds the baseline level and may serve to restore the functional state of the repeatedly stimulated cortical areas. Here we investigated whether the changes in MEG-based parameters previously associated with inhibition (increases in periodic alpha-beta power, increases in alpha frequency, and steepness of aperiodic 1/f slope) were sensitive to post-stimulus enhancement of inhibition compared to the resting state. We also investigated whether changes in these parameters are proportional to the intensity of the preceding visual stimulation. We then tested whether changes in MEG parameters associated with post-stimulus inhibition correlated with subjective sensory sensitivity and avoidance.

Our main results show that all four investigated parameters depend on visual stimulation intensity. Their intensity-dependent changes concordantly indicate a greater shift to functional inhibition after cessation of more intense versus less intense visual input, and significant correlations between some of these parameters (and their changes) suggest that they are likely to share some common underlying mechanisms.

In our study, the aperiodic slope demonstrated changes consistent with those of the other MEG parameters only when it was measured in the high-frequency range, but not in the broad range of frequencies. Therefore, in the following discussion, we focus on the high-frequency slope and discuss the difference between the two methods of aperiodic slope estimation in a separate methodological section.

### MEG indices of post-stimulus functional inhibition scale in proportion to the intensity of visual stimulation

We found that all four examined MEG indices were sensitive to the strength of preceding visual stimulation (faster versus slower drift rate of visual gratings) and, except for the PAF, displayed correlated changes that were compatible with an intensity-dependent increase in post-stimulus functional inhibition. Note, that the term ‘post-stimulus’ here refers to the time window after visual stimulation has ceased. These results support the hypothesis that post-stimulus inhibition increases proportionally to the strength of preceding excitation (Haigh et al., 2015; Mullinger et al., 2017). Whereas associations between the intensity of preceding visual stimulation and increases in alpha and/or beta power in the post-stimulus interval have been reported previously (Mullinger et al., 2017; Stevenson et al., 2011), we showed such an association for PAF and aperiodic 1/f slope for the first time.

Greater intensity of the preceding stimulation was associated with a steeper aperiodic slope in stimulated visual areas (Figure 3). Since the steepness of the aperiodic slope measured in high-frequency range of the power spectrum (>30 Hz) is associated with an overall decrease in neural excitability (Gao et al., 2017; Wiest et al., 2023), this finding suggests that greater activation of the visual cortex by incoming visual input is followed by a proportionally greater functional inhibition in the post-stimulus interval.

Consistent with the results of Muthukumaraswamy and Liley (Muthukumaraswamy & Liley, 2018), who analyzed resting state MEG and EEG, the greater negativity of the aperiodic 1/f slope estimated at posterior sensors correlates with greater power of periodic alpha-beta oscillations. Here, this correlation was observed for both resting state and post-stimulus intervals (Table 2). Moreover, changes in these parameters as a function of stimulation intensity (from ‘after-slow’ to ‘after-fast’ interval) were also correlated: greater negativity (steepness) of the slope was associated with a greater increase in the power of periodic alpha-beta oscillations (Table 3b). Thus, the strength and direction of intensity-related changes in aperiodic 1/f slope and power of periodic alpha-beta oscillations point in the same direction: once the intensity of the visual stimulation is going up, the post-stimulus state of the visual cortex undergoes a shift towards inhibition. This push-pull relationship between the intensity of the visual input and post-stimulus changes in inhibition-sensitive MEG parameters is compatible with the hypothesis that functional inhibition plays a homeostatic role in balancing cortical excitability after intensive stimulation (Mullinger et al., 2017).

Weak but significant correlations between changes in post-stimulus periodic power and aperiodic slope suggest some similarity in the underlying mechanisms. This is not unexpected, since a number of studies relate steeper aperiodic slope and higher alpha-beta power to a state of increased functional inhibition characterized by reduction of neuronal firing (alpha-beta power: (Bonnefond & Jensen, 2015; Haegens et al., 2011; Iemi et al., 2022; van Kerkoerle et al., 2014); aperiodic slope: (Gao et al., 2017; Wiest et al., 2023))

While, according to modeling, animal and intracranial studies the link between changes in aperiodic slope and neural excitability is straightforward – a steeper slope directly reflects stronger local inhibition (Gao et al., 2017; Wiest et al., 2023), the associations between inhibition and changes in alpha-beta power are more complex. Functionally relevant increase of alpha-beta power is thought to reflect either inhibitory top-down control (Bonnefond & Jensen, 2012; Jensen, Gelfand, Kounios, & Lisman, 2002; Klimesch, Doppelmayr, Schwaiger, Auinger, & Winkler, 1999; Payne, Guillory, & Sekuler, 2013), or be secondary to competitive interactions between strongly activated task-relevant cortical regions and task-irrelevant ones, which neuronal activity is inhibited to prevent interference (Jensen, 2023; Noonan, Crittenden, Jensen, & Stokes, 2018). At first glance, the latter interpretation is not consistent with the current finding, because the intensity-dependent post-stimulus increase in alpha (beta) power is localized to the task-relevant visual cortical areas. However, post-stimulus alpha-beta synchronization may be a delayed consequence of competitive interactions between populations of neurons in the same cortical area occuring during sensory stimulation (i.e. surround inhibition). For example, Boorman et al have found that the regions surrounding the stimulated receptive field in somatosensory cortex in animals demonstrated a negative hemodynamic response that outlasted the ‘positive hemodynamic response’ and correlated with prolonged (300-2000 ms after the termination of the stimulation) suppression of broad-band gamma power (i.e., suppression of neural activity, see (Manning, Jacobs, Fried, & Kahana, 2009)). Since large high-contrast moving gratings used in our study activate both neurons’ receptive fields and their suppressive surrounds and induce strong surround inhibition (Gieselmann & Thiele, 2008), alpha-beta synchronization and steepening of the 1/f aperiodic slope that we observed in the post-stimulus interval may reflect such local residual inhibitory processes resulting from competitive interactions *within the same visual area.* This hypothesis is indirectly supported by the results of the iEEG/fMRI study of Harvey et al (Harvey et al., 2013) who showed that surround inhibition in the human primary visual cortex is associated with local increases in alpha activity in the population receptive fields. Presumably, cessation of stimulation can trigger synchronization of these local alpha oscillators over an extensive region of visual cortex, resulting in alpha-beta ‘rebound’ detected by MEG. In this case, the dependence of the post-stimulus alpha-beta increase on the grating drift rate (Figure 3) can be explained by stronger surround inhibition induced by stimuli drifting at a higher than at a lower rate (see (Orekhova et al., 2020) for discussion). Thus, the post-stimulus alpha-beta ‘rebound’ might be a secondary consequence of excitatory-inhibitory interactions driven in the feed-forward way by preceding visual input.

On the other hand, we found some indirect evidence for top-down influences on post-stimulus periodic alpha-beta power. In our participants, subjective hypersensitivity estimated by Low Neurological Threshold score correlated with а relatively *greater* intensity-related enhancement in alpha-beta power in the post-stimulus interval (Figure 7, Table 4). Notably, this correlation was absent for the aperiodic slope (Table 4). This suggests that some additional factors upregulate post-stimulus alpha-beta power over and above the local residual inhibitory processes. According to the previous literature, periodic alpha power in sensory cortical areas, although correlate with the strength of high-frequency neural response to sensory input - a direct measure of cortical excitability (Iemi et al., 2022), is also modulated by top-down processes, such as e.g., attention engagement/disengagement that are accessible to awareness providing a basis for subjective feeling of stimulation intensity (Kerr et al., 2011; Klimesch, 2012; Peng, Babiloni, Mao, & Hu, 2015; Samaha, Iemi, Haegens, & Busch, 2020).

It is likely that, similar to people with somatic anxiety (Bouziane et al., 2022), healthy individuals scoring high on the ‘Low Neurological Threshold’ exerted greater top-down inhibitory control over excitability of the exteroceptive sensory (in this case visual) cortical areas than less sensitive individuals. Stronger top-down inhibitory control over the highly excited visual cortex may lead to a proportionally higher increase of alpha-beta power after intensive stimulation and explains its link to subjective sensory sensitivity and avoidance.

These considerations suggest that post-stimulus alpha-beta ‘rebound’ in visual cortex may reflect both feed-forward and feedback inhibitory processes, and evaluation of their relative contributions to this phenomenon deserves future studies (see, e.g., (Khan et al., 2015)).

### Dissociation between changes in periodic alpha-beta power and 1/f aperiodic slope from resting state to post-stimulus interval

The contrast between the resting state and post-stimulus period showed that the latter condition was characterized by steepening of the aperiodic slope in the visual cortex (Figures 3, 5). This finding, along with the increase in periodic beta power and PAF in posterior sensors (Figure 3), is consistent with an increase of inhibition during post-stimulus period compared to rest. Since participants were presented with the same fixation cross in both conditions, these results cannot be explained by differences in visual input but rather reflect internal processes related to the cessation of visual stimulation. Thus, these MEG findings agree with the results of other MEG/EEG studies in humans (Mullinger et al., 2017; Mullinger et al., 2013; Stevenson et al., 2011) and neuronal studies in animals (Boorman et al., 2015; Logothetis et al., 2001; Shmuel et al., 2006), in which a post-stimulus enhancement of inhibition above the baseline level was observed.

However, the pattern of differences in MEG parameters between the post-stimulus period and rest cannot be explained solely by post-stimulus increase of inhibition in the stimulated cortical areas. First, cortical topography of condition-related changes was markedly different for aperiodic slope and beta power (Figure 5). Whereas the difference in the aperiodic slope between rest and post-stimulus interval was limited to the visual cortex and surrounding areas, the differences in beta power were widespread.

One obvious source of increased beta power during the post-stimulus period relative to rest in our study is the post-movement beta rebound (PMBR) associated with the button press that subjects performed in the visual task. Since the PMBR begins about 230 ms after movement termination and lasts several hundred milliseconds (Jurkiewicz, Gaetz, Bostan, & Cheyne, 2006), it should coincide with our analysis window (Figure 1B). However, the maximal amplitude of PMBR is expected in the precentral gyrus, contralateral to the movement side (Gaetz, Macdonald, Cheyne, & Snead, 2010; Jurkiewicz et al., 2006; Wilson et al., 2010). Therefore, the increase of beta power in large-scale cortical networks in the post-stimulus period (Figure 5) cannot be accounted for solely by PMBR.

Interestingly, the most pronounced relative increase in beta power during post-stimulus period compared to rest was observed in cingulate cortex, orbitofrontal cortex and insula. Since these areas are involved in emotional and autonomic regulation (Ferraro et al., 2022; Hänsel & von Känel, 2008; Stevens, Hurley, & Taber, 2011), the beta synchronization may be related to the subjective experience of visual stimulation. While moving visual gratings used in our study were not explicitly aversive, subjects often rated them as unpleasant (Orekhova et al., 2018). Therefore, the post-stimulus beta power increase in the ventromedial prefrontal cortex and anterior insula may reflect downregulation of emotional arousal and/or autonomic activity during our ‘mildly unpleasant’ visual experiment. The lack of influence of intensity of the preceding stimulation on post-stimulus beta power in the respective high-order cortical areas (Figure 6) can be explained by the tonic nature of these regulatory processes: stimuli were presented at random, and the stimulation intensity (drift rate) in the upcoming trial could not be predicted.

Although the mechanisms of the widespread increase in beta oscillations after cessation of intense visual stimulation is beyond the scope of the present work, we can cautiously offer an explanation. The increase in ongoing high-frequency alpha and/or beta power in the ventromedial prefrontal cortex has been previously observed after vigorous physical activity (Hosang, Mouchlianitis, Guérin, & Karageorghis, 2022) and linked to the enhanced blood level of serotonin (5-HT) (Fumoto et al., 2010). The 5-HT is known to attenuate the brain excitatory state induced by physiological arousal due to its inhibitory effect on cortical neuronal activity through 5-HT1A receptors, which are abundant in the ventromedial prefrontal cortex (Celada, Puig, & Artigas, 2013). A recent study reported that a stressful event, electrical stimulation of a finger, elicits a post-stimulus increase of MEG beta power in the anterior cingulum that strongly correlates with transient short-latency inhibition of peripheral sympathetic nerve activity, suggesting a common mechanism underlying both responses (Riaz et al., 2022). Given that 5 HT 1A receptors are implicated in both sympatho-inhibition (Jonnakuty & Gragnoli, 2008) and down-regulation of neuronal activity in ventromedial PFC (Celada et al., 2013), a serotonin-dependent pathway is a possible candidate mechanism for inhibitory control of post-stressful cortical inactivation through beta synchronization. A similar mechanism may work to increase beta power and tonically attenuate the excitatory state associated with unpleasant visual stimulation in our study. Regardless of the exact mechanisms, our results show that reduced excitability of the visual areas does not fully characterize inhibition-related differences in beta power between rest and post-stimulus intervals.

Furthermore, we found no differences in alpha power between rest and post-stimulus conditions. Thus, post-stimulus alpha power in the visual areas is reliably modulated by the intensity of visual stimulation, but not by the very fact of the presence of such stimulation compared to its absence (compare Figures 3A and 3B). This finding is broadly consistent with results of Mullinger and colleagues (Mullinger et al., 2017), who found little or no changes in alpha power between resting state and intervals following flickering of static visual gratings, but a highly significant increase in alpha power following disappearance of flickering compared to static visual gratings.

A likely explanation of these findings is the high inter-subject variability of physiological arousal in a poorly controlled resting state. Since low level of physiological arousal is associated with high resting alpha power (Barry, De Blasio, Fogarty, & Clarke, 2020; Hutt & Lefebvre, 2022), the transition from a resting state to a more active state during a visual task may lead to a decrease in alpha power not only during visual stimulation, but also in the inter-stimulus intervals. Indeed, we found that this was the case in our participants with higher-than-average alpha power at rest. As the increase in the tonic arousal and post-stimulus inhibition counteract each other, their cumulative effect may result in the absence of difference in alpha power between rest and post-stimulus condition that we observed at the group level.

To sum up, the contrast between the time intervals following cessation of intensive visual stimulation and the resting state revealed partly dissimilar changes in the studied MEG parameters, which could not be explained exclusively by post-stimulus functional inhibition of the stimulated visual areas. The discrepancy between changes in different inhibition-sensitive MEG parameters may be explained by differences in vigilance/tonic arousal between post-stimulus condition and the resting state.

### Peak alpha frequency does not correlate with other MEG parameters sensitive to inhibition and changes during menstrual cycle

According to interpretation proposed by Babu Henry Samuel, 2018 et al (Babu Henry Samuel et al., 2018), the post-stimulus and intensity-related increase in PAF (Figure 2) suggest increased inhibition. Thus, the changes in PAF were in line with changes in other MEG parameters studied (alpha-beta power, aperiodic slope). However, changes in PAF did not correlate with changes in other ‘inhibition-based’ MEG parameters, suggesting that PAF provides distinct information on post-stimulus functional inhibition (Tables 3A,B).

The lack of correlations between PAF and alpha power agrees with the previous literature, which shows that, depending on the experimental condition, an increase in alpha frequency may be accompanied by either decrease (Haegens, Cousijn, Wallis, Harrison, & Nobre, 2014) (for review see (Mierau, Klimesch, & Lefebvre, 2017)) or increase ((Wianda & Ross, 2019); the present study) in alpha power. It is also in line with our observation that only PAF was modulated by the phase of the MC. Regardless of the experimental condition, PAF was higher in the luteal phase than in the follicular phase. The same differences in the alpha frequency between luteal and follicular phases during resting state have been previously described in several studies (Bazanova, Kondratenko, Kuz’minova, Muravleva, & Petrova, 2014; Brötzner, Klimesch, Doppelmayr, Zauner, & Kerschbaum, 2014; Haraguchi et al., 2021). In general, functional dissociation between frequency and power of alpha oscillations is inconsistent with unitary explanation of changes in alpha frequency and power through the single mechanism (Peterson & B., 2017).

An interesting parallel between PAF and gamma oscillations frequency arises here. Experimental research (Haegens et al., 2011; Mann & Mody, 2010; Walker & Semyanov, 2008) and some modeling studies (Vierling-Claassen, Cardin, Moore, & Jones, 2010) posit that frequency of both gamma and alpha oscillations is determined by the pace of rhythmic inhibition of neuronal spiking, which in turn depends on excitability of inhibitory neurons, albeit of different types (Vierling-Claassen et al., 2010). Indeed, modulation of the tonic excitability of inhibitory interneurons in mice dramatically affects the frequency (but not amplitude) of gamma oscillations (Mann & Mody, 2010). Continuing this analogy, animal studies have shown that natural (e.g., during pregnancy) or experimental changes in steroid hormone concentrations alter gamma frequency, most likely through complex modulation of tonic excitability of inhibitory neurons (Ferando & Mody, 2013). There is an intriguing possibility that MC phase-related change in PAF and correlation of PAF with progesterone concentration (Bazanova et al., 2014) are also caused by modulation of inhibitory neurons’ excitability by steroid hormones. Since excitability of inhibitory neurons is only one of many factors regulating E/I balance, this may explain the concordant (indicating increased post-stimulus functional inhibition with increasing strength of prior input) but uncorrelated changes in PAF and other MEG parameters that, unlike PAF, index the resulting changes in E/I ratio.

Overall, our results suggest the mechanisms mediating changes in PAF are different from those underlying changes in alpha-beta power and aperiodic slope. Although the current explanation for these mechanisms is speculative, the functional dissociation we found between PAF and other putative measures of E/I ratio deserves further investigation because of its physiological relevance and potential implications for the use of MEG/EEG indexes of functional inhibition in translational studies.

### Methodological aspects of spectral slope estimation

The aperiodic 1/f-like slope of the power spectrum, estimated in the range 35-45 Hz, exhibited functionally meaningful modulations compatible with post-stimulus increase in functional inhibition (Figures 3A,B). In contrast, the spectral slope estimated in 2-40 Hz range using FOOOF (Donoghue et al., 2020) did not change across conditions. Additionally, the previously reported negative correlation between periodic alpha power and the aperiodic slope (Muthukumaraswamy & Liley, 2018; Podvalny et al., 2015) was reproduced in our study only for the slope estimated in the high-frequency range. The correlation between spectral slope and periodic alpha-beta power is expected because both these parameters correlate with spiking activity (Chapeton et al., 2019; Dougherty et al., 2017; Gao et al., 2017; Haegens et al., 2011; Wiest et al., 2023). Remarkably, we found no significant correlations between the slopes estimated in two different ways.

In the literature, the aperiodic slope is often estimated either over a wide range of frequencies after separating between periodic and aperiodic parts of the power spectrum, or in the high-frequency range (>30 Hz), where periodic activity is usually absent, and direct comparison between these methods is rarely performed (see (Lendner et al., 2020; Maschke et al., 2023) for an exception). However, both of these methods have limitations.

Reliable estimation of the aperiodic slope in the high-frequency range can be hampered by the presence of myogenic artifacts and/or sensor noise. For example, Maschke et al (Maschke et al., 2023) reported that the steepness of the EEG spectral slope estimated in the 30-45 Hz range (but not in the 1-45 Hz range) predicted the response to propofol-induced anesthesia and the participant’s level of consciousness before the anesthetic state but did not rule out that this result was due to myogenic artifacts. In our study, condition- and intensity-related changes in the 35-45 Hz slope were unlikely to be explained by differences in muscular activity. Although myogenic artifacts may propagate to some extent across MEG sensors, they predominantly affect peripheral sensors located over active muscles (Goncharova, McFarland, Vaughan, & Wolpaw, 2003). In contrast, we found that differences in the 35-45 Hz aperiodic slope between conditions disappeared for posterior gradiometers most contaminated by myogenic activity (Supplementary Results). This finding strongly supports our hypothesis of a neural origin of condition- and intensity-related modulations of the 35-45 Hz aperiodic slope.

Estimating the aperiodic slope based on broadband activity has other limitations (see (Gerster et al., 2022) for review). A recent MEG study showed that throughout the cerebral cortex, the aperiodic part of PSD is best approximated by a model with *two* slopes separated by a knee at 15 Hz (Ibarra Chaoul & Siegel, 2021). Therefore, FOOOF algorithm, which describes the aperiodic power of human brain activity by a single power law, may not be optimal for slope estimation. Since our experimental design (1 s epochs) did not allow the IRASA approach used by Ibarra Chaoul and colleagues (Ibarra Chaoul & Siegel, 2021), we attempted to fit MEG spectra using FOOOF model with a ‘knee’ mode. This approach, however, resulted in highly variable knee frequencies and occasionally unrealistically steep slopes (see Supplementary Results).

As the previous animal and modeling study have shown the link between neural excitability and the aperiodic 1/f slope at high frequencies (>30 Hz) (Gao et al., 2017), we believe that the spectral slope we estimated in the 35-45 Hz range, where rhythmic activity is absent, provides a more realistic approximation of 1/f slope of aperiodic neural activity than the slope estimated in the broad frequency range using FOOOF. In addition to its functional relevance, the 35-45 Hz slope is a sufficiently reliable measure of inter-individual variability, as it shows high rank-order stability between the two measurements separated by several days or weeks (Table 1). This is consistent with our previous results showing that aperiodic slope estimated in the frequency range 35-45 Hz predicts child age and differentiates between autistic children with below-average and normal IQ (V.O. Manyukhina et al., 2022).

The aperiodic slope becomes an increasingly popular index of the E/I ratio in neuropsychiatric and neurodevelopmental disorders, such as schizophrenia (Molina et al., 2020), attention deficit hyperactivity disorders (Arnett, Rutter, & Stein, 2022; Ostlund, Alperin, Drew, & Karalunas, 2021; Robertson et al., 2019), obsessive-compulsive disorder (Perera, Mallawaarachchi, Bailey, Murphy, & Fitzgerald, 2023), depression (Veerakumar et al., 2019), Rett syndrome (Roche et al., 2019), Fragile X syndrome (Wilkinson & Nelson, 2021) and autism spectrum disorders (Carter Leno et al., 2022). Many of these recent studies used EEG and estimated a single aperiodic slope in a broad frequency range (e.g. 1-40 Hz). Although PSD scaling at frequencies below 10 Hz differs in MEG and EEG (Dehghani, Bédard, Cash, Halgren, & Destexhe, 2010), the presence of a ‘knee’ may not be specific to MEG. Indeed, intracranial EEG studies show that the slope of the aperiodic part of the power spectrum is steeper at frequencies above than below ∼10 Hz (see Figure 3A in (Sheehan, Sreekumar, Inati, & Zaghloul, 2018)). Some anesthetics differently affect aperiodic spectral slopes at higher and lower frequencies in surface EEG (Colombo et al., 2019), which suggests that the factors contributing to the aperiodic activity may differ in different frequency ranges. Moreover, aperiodic slopes in the high and low frequency range are differently affected by visual deprivation. A significantly steeper (more negative) low-frequency (1.5 to 19.5 Hz) and significantly higher (less negative) high-frequency (20 to 45 Hz) slopes were found in blind patients and patients who underwent late cataract removal than in normally seeing individuals (Ossandón et al., 2023). Collectively, these and our present findings urge caution while interpreting the aperiodic slope estimated over a wide range of frequencies.

## Limitations

Our study has several limitations. First, the short 1-second post-stimulus interval we used (0.2 s – 1.2 s after stimulus cessation) may have missed a portion of the post-stimulus inhibitory response, which in MEG/EEG may last several seconds after stimulation stopped (Mullinger et al., 2017; Stevenson et al., 2011). Longer inter-stimulus intervals would allow a more detailed analysis of the post-stimulus changes in inhibition. Second, our study included a relatively small sample of healthy adult women of reproductive age. A study on men is warranted before our findings can be extended to the general population. Another limitation is the use of the FOOOF algorithm to isolate periodic activity, since this method was not optimal for separating the periodic and aperiodic parts of the power spectra in our study. However, since the spectral aperiodic part of the spectrum estimated with FOOOF did not vary significantly as a function of the experimental condition, we hope that its application does not systematically affect the estimation of changes in periodic activity.

## Conclusions

Our results confirm the previous finding linking the ‘overshoot’ of alpha-beta power following cessation of visual stimulation to functional inhibition and extend them by showing that the 35-45 Hz aperiodic 1/f slope and alpha peak frequency also reflect a shift toward inhibition in the post-stimulus activity of the human visual cortex. The concordant changes in all inhibition-sensitive parameters associated with changes in stimulation intensity are consistent with the idea that post-stimulus inhibition is proportional to the intensity of the preceding visual stimulation. Whereas changes in the 35-45 Hz aperiodic slope from resting to post-stimulus period clearly indicate enhanced inhibition in the stimulated visual cortex, the concurrent modulations of periodic alpha and beta power are more complex and presumably reflect processes not limited to the down-regulation of the E/I ratio in stimulated sensory cortical areas. Finally, the aperiodic 1/f slope showed the expected behavior, consistent with previous findings, only when it was estimated in the high-frequency range. This result calls for caution when interpreting the aperiodic slope estimated over a wide range of frequencies.

## Data and code availability

The data and code used to derive the results of this study are available upon request from the corresponding author.

## Author contributions

*Viktoriya O. Manyukhina*: Performed experiments, Analyzed data, Interpreted results of experiments, Prepared figures, Drafted manuscript, Edited and revised manuscript, Approved final version of manuscript.

*Andrey O. Prokofyev*: Conceived and designed research, Performed experiments.

*Tatiana S. Obukhova*: Performed experiments.

*Tatiana A. Stroganova*: Conceived and designed research, Interpreted results of experiments, Edited and revised manuscript, Approved final version of manuscript.

*Elena V. Orekhova*: Conceived and designed research, Analyzed data, Interpreted results of experiments, Prepared figures, Drafted manuscript, Edited and revised manuscript, Approved final version of manuscript.

## Acknowledgments

We sincerely thank all of the participants who participated in this study. The research was conducted within the framework of the state assignment of the Ministry of Education of the Russian Federation from 13.02.2023 (project N 073-00038-23-02). Part of the study (comparison of different methods for estimating aperiodic spectral slopes) was supported by the Russian Science Foundation (project N 22-25-00419).

## References

Ahmad, J., Ellis, C., Leech, R., Voytek, B., Garces, P., Jones, E., … McAlonan, G. (2022). From mechanisms to markers: novel noninvasive EEG proxy markers of the neural excitation and inhibition system in humans. Transl Psychiatry, 12(1), 467. doi:10.1038/s41398-022-02218-z

Anticevic, A., & Murray, J. D. (2017). Rebalancing Altered Computations: Considering the Role of Neural Excitation and Inhibition Balance Across the Psychiatric Spectrum. Biol Psychiatry, 81(10), 816–817. doi:10.1016/j.biopsych.2017.03.019

Arnett, A. B., Rutter, T. M., & Stein, M. A. (2022). Neural Markers of Methylphenidate Response in Children With Attention Deficit Hyperactivity Disorder. Front Behav Neurosci, 16, 887622. doi:10.3389/fnbeh.2022.887622

Babu Henry Samuel, I., Wang, C., Hu, Z., & Ding, M. (2018). The frequency of alpha oscillations: Task-dependent modulation and its functional significance. Neuroimage, 183, 897–906. doi:10.1016/j.neuroimage.2018.08.063

Bargary, G., Furlan, M., Raynham, P. J., Barbur, J. L., & Smith, A. T. (2015). Cortical hyperexcitability and sensitivity to discomfort glare. Neuropsychologia, 69, 194–200. doi:10.1016/j.neuropsychologia.2015.02.006

Barry, R. J., De Blasio, F. M., Fogarty, J. S., & Clarke, A. R. (2020). Natural alpha frequency components in resting EEG and their relation to arousal. Clin Neurophysiol, 131(1), 205–212. doi:10.1016/j.clinph.2019.10.018

Bazanova, O. M., Kondratenko, A. V., Kuz’minova, O. I., Muravleva, K. B., & Petrova, S. E. (2014). [EEG alpha indices in dependence on the menstrual cycle phase and salivary progesterone]. Fiziol Cheloveka, 40(2), 31–40.

Bonnefond, M., & Jensen, O. (2012). Alpha oscillations serve to protect working memory maintenance against anticipated distracters. Curr Biol, 22(20), 1969–1974. doi:10.1016/j.cub.2012.08.029

Bonnefond, M., & Jensen, O. (2015). Gamma activity coupled to alpha phase as a mechanism for top-down controlled gating. PLoS One, 10(6), e0128667. doi:10.1371/journal.pone.0128667

Boorman, L., Harris, S., Bruyns-Haylett, M., Kennerley, A., Zheng, Y., Martin, C., … Berwick, J. (2015). Long-latency reductions in gamma power predict hemodynamic changes that underlie the negative BOLD signal. J Neurosci, 35(11), 4641–4656. doi:10.1523/jneurosci.2339-14.2015

Bouziane, I., Das, M., Friston, K. J., Caballero-Gaudes, C., & Ray, D. (2022). Enhanced top-down sensorimotor processing in somatic anxiety. Transl Psychiatry, 12(1), 295. doi:10.1038/s41398-022-02061-2

Brötzner, C. P., Klimesch, W., Doppelmayr, M., Zauner, A., & Kerschbaum, H. H. (2014). Resting state alpha frequency is associated with menstrual cycle phase, estradiol and use of oral contraceptives. Brain Res, 1577(100), 36–44. doi:10.1016/j.brainres.2014.06.034

Brown, C. E., & Dunn, W. (2002). Adolescent/adult sensory profile: User’s manual. A San Antonio, TX: The Psychological Corporation.

Carter Leno, V., Begum-Ali, J., Goodwin, A., Mason, L., Pasco, G., Pickles, A., … Jones, E. J. H. (2022). Infant excitation/inhibition balance interacts with executive attention to predict autistic traits in childhood. Mol Autism, 13(1), 46. doi:10.1186/s13229-022-00526-1

Celada, P., Puig, M. V., & Artigas, F. (2013). Serotonin modulation of cortical neurons and networks. Front Integr Neurosci, 7, 25. doi:10.3389/fnint.2013.00025

Chapeton, J. I., Haque, R., Wittig, J. H., Jr., Inati, S. K., & Zaghloul, K. A. (2019). Large-Scale Communication in the Human Brain Is Rhythmically Modulated through Alpha Coherence. Curr Biol, 29(17), 2801–2811.e2805. doi:10.1016/j.cub.2019.07.014

Chen, J. J., & Pike, G. B. (2009). Origins of the BOLD post-stimulus undershoot. Neuroimage, 46(3), 559–568. doi:10.1016/j.neuroimage.2009.03.015

Chen, Q., Deister, C. A., Gao, X., Guo, B., Lynn-Jones, T., Chen, N., … Feng, G. (2020). Dysfunction of cortical GABAergic neurons leads to sensory hyper-reactivity in a Shank3 mouse model of ASD. Nat Neurosci, 23(4), 520–532. doi:10.1038/s41593-020-0598-6

Colombo, M. A., Napolitani, M., Boly, M., Gosseries, O., Casarotto, S., Rosanova, M., Sarasso, S. (2019). The spectral exponent of the resting EEG indexes the presence of consciousness during unresponsiveness induced by propofol, xenon, and ketamine. Neuroimage, 189, 631–644. doi:10.1016/j.neuroimage.2019.01.024

Dehghani, N., Bédard, C., Cash, S. S., Halgren, E., & Destexhe, A. (2010). Comparative power spectral analysis of simultaneous elecroencephalographic and magnetoencephalographic recordings in humans suggests non-resistive extracellular media. J Comput Neurosci, 29(3), 405–421. doi:10.1007/s10827-010-0263-2

Donoghue, T., Haller, M., Peterson, E. J., Varma, P., Sebastian, P., Gao, R., … Voytek, B. (2020). Parameterizing neural power spectra into periodic and aperiodic components. Nat Neurosci, 23(12), 1655–1665. doi:10.1038/s41593-020-00744-x

Dougherty, K., Cox, M. A., Ninomiya, T., Leopold, D. A., & Maier, A. (2017). Ongoing Alpha Activity in V1 Regulates Visually Driven Spiking Responses. Cereb Cortex, 27(2), 1113–1124. doi:10.1093/cercor/bhv304

Ethridge, L. E., White, S. P., Mosconi, M. W., Wang, J., Pedapati, E. V., Erickson, C. A., … Sweeney, J. A. (2017). Neural synchronization deficits linked to cortical hyper-excitability and auditory hypersensitivity in fragile X syndrome. Mol Autism, 8, 22. doi:10.1186/s13229-017-0140-1

Ferando, I., & Mody, I. (2013). Altered gamma oscillations during pregnancy through loss of δ subunit-containing GABA(A) receptors on parvalbumin interneurons. Front Neural Circuits, 7, 144. doi:10.3389/fncir.2013.00144

Ferraro, S., Klugah-Brown, B., Tench, C. R., Bazinet, V., Bore, M. C., Nigri, A., … Becker, B. (2022). The central autonomic system revisited - Convergent evidence for a regulatory role of the insular and midcingulate cortex from neuroimaging meta-analyses. Neurosci Biobehav Rev, 142, 104915. doi:10.1016/j.neubiorev.2022.104915

Fischl, B., Salat, D. H., Busa, E., Albert, M., Dieterich, M., Haselgrove, C., … Dale, A. M. (2002). Whole brain segmentation: Automated labeling of neuroanatomical structures in the human brain. Neuron, 33(3), 341–355. doi:10.1016/S0896-6273(02)00569-X

Fumoto, M., Oshima, T., Kamiya, K., Kikuchi, H., Seki, Y., Nakatani, Y., … Arita, H. (2010). Ventral prefrontal cortex and serotonergic system activation during pedaling exercise induces negative mood improvement and increased alpha band in EEG. Behav Brain Res, 213(1), 1–9. doi:10.1016/j.bbr.2010.04.017

Gaetz, W., Macdonald, M., Cheyne, D., & Snead, O. C. (2010). Neuromagnetic imaging of movement-related cortical oscillations in children and adults: age predicts post-movement beta rebound. Neuroimage, 51(2), 792–807. doi:10.1016/j.neuroimage.2010.01.077

Gao, R., Peterson, E. J., & Voytek, B. (2017). Inferring synaptic excitation/inhibition balance from field potentials. Neuroimage, 158, 70–78. doi:10.1016/j.neuroimage.2017.06.078

Gerster, M., Waterstraat, G., Litvak, V., Lehnertz, K., Schnitzler, A., Florin, E., … Nikulin, V. (2022). Separating Neural Oscillations from Aperiodic 1/f Activity: Challenges and Recommendations. Neuroinformatics, 20(4), 991–1012. doi:10.1007/s12021-022-09581-8

Gieselmann, M. A., & Thiele, A. (2008). Comparison of spatial integration and surround suppression characteristics in spiking activity and the local field potential in macaque V1. Eur J Neurosci, 28(3), 447–459. doi:10.1111/j.1460-9568.2008.06358.x

Goncharova, II, McFarland, D. J., Vaughan, T. M., & Wolpaw, J. R. (2003). EMG contamination of EEG: spectral and topographical characteristics. Clin Neurophysiol, 114(9), 1580–1593. doi:10.1016/s1388-2457(03)00093-2

Gramfort, A., Luessi, M., Larson, E., Engemann, D. A., Strohmeier, D., Brodbeck, C., … Hamalainen, M. (2013). MEG and EEG data analysis with MNE-Python. Frontiers in Neuroscience, 7. doi:10.3389/fnins.2013.00267

Haegens, S., Cousijn, H., Wallis, G., Harrison, P. J., & Nobre, A. C. (2014). Inter- and intra-individual variability in alpha peak frequency. Neuroimage, 92(100), 46–55. doi:10.1016/j.neuroimage.2014.01.049

Haegens, S., Nácher, V., Luna, R., Romo, R., & Jensen, O. (2011). α-Oscillations in the monkey sensorimotor network influence discrimination performance by rhythmical inhibition of neuronal spiking. Proc Natl Acad Sci U S A, 108(48), 19377–19382.

Haigh, S. M., Cooper, N. R., & Wilkins, A. J. (2015). Cortical excitability and the shape of the haemodynamic response. Neuroimage, 111, 379–384. doi:10.1016/j.neuroimage.2015.02.034

Hänsel, A., & von Känel, R. (2008). The ventro-medial prefrontal cortex: a major link between the autonomic nervous system, regulation of emotion, and stress reactivity? Biopsychosoc Med, 2, 21. doi:10.1186/1751-0759-2-21

Haraguchi, R., Hoshi, H., Ichikawa, S., Hanyu, M., Nakamura, K., Fukasawa, K., … Shigihara, Y. (2021). The Menstrual Cycle Alters Resting-State Cortical Activity: A Magnetoencephalography Study. Front Hum Neurosci, 15, 652789. doi:10.3389/fnhum.2021.652789

Harvey, B. M., Vansteensel, M. J., Ferrier, C. H., Petridou, N., Zuiderbaan, W., Aarnoutse, E. J., … Dumoulin, S. O. (2013). Frequency specific spatial interactions in human electrocorticography: V1 alpha oscillations reflect surround suppression. Neuroimage, 65, 424–432. doi:10.1016/j.neuroimage.2012.10.020

Havlicek, M., Roebroeck, A., Friston, K., Gardumi, A., Ivanov, D., & Uludag, K. (2015). Physiologically informed dynamic causal modeling of fMRI data. Neuroimage, 122, 355–372. doi:10.1016/j.neuroimage.2015.07.078

Hosang, L., Mouchlianitis, E., Guérin, S. M. R., & Karageorghis, C. I. (2022). Effects of exercise on electroencephalography-recorded neural oscillations: a systematic review. International Review of Sport and Exercise Psychology, 1–54. doi:10.1080/1750984X.2022.2103841

Hutt, A., & Lefebvre, J. (2022). Arousal Fluctuations Govern Oscillatory Transitions Between Dominant [Formula: see text] and [Formula: see text] Occipital Activity During Eyes Open/Closed Conditions. Brain Topogr, 35(1), 108–120. doi:10.1007/s10548-021-00855-z

Ibarra Chaoul, A., & Siegel, M. (2021). Cortical correlation structure of aperiodic neuronal population activity. Neuroimage, 245, 118672. doi:10.1016/j.neuroimage.2021.118672

Iemi, L., Gwilliams, L., Samaha, J., Auksztulewicz, R., Cycowicz, Y. M., King, J. R., … Haegens, S. (2022). Ongoing neural oscillations influence behavior and sensory representations by suppressing neuronal excitability. Neuroimage, 247, 118746. doi:10.1016/j.neuroimage.2021.118746

Jensen, O. (2023). Gating by alpha band inhibition revised: a case for a secondary control mechanism. PsyArXiv. doi:10.31234/osf.io/7bk32

Jensen, O., Gelfand, J., Kounios, J., & Lisman, J. E. (2002). Oscillations in the alpha band (9-12 Hz) increase with memory load during retention in a short-term memory task. Cereb Cortex, 12(8), 877–882. doi:10.1093/cercor/12.8.877

Jonnakuty, C., & Gragnoli, C. (2008). What do we know about serotonin? J Cell Physiol, 217(2), 301–306. doi:10.1002/jcp.21533

Jurkiewicz, M. T., Gaetz, W. C., Bostan, A. C., & Cheyne, D. (2006). Post-movement beta rebound is generated in motor cortex: evidence from neuromagnetic recordings. Neuroimage, 32(3), 1281–1289. doi:10.1016/j.neuroimage.2006.06.005

Kerr, C. E., Jones, S. R., Wan, Q., Pritchett, D. L., Wasserman, R. H., Wexler, A., … Moore, C. I. (2011). Effects of mindfulness meditation training on anticipatory alpha modulation in primary somatosensory cortex. Brain Res Bull, 85(3-4), 96–103. doi:10.1016/j.brainresbull.2011.03.026

Khan, S., Hashmi, J. A., Mamashli, F., Michmizos, K., Kitzbichler, M. G., Bharadwaj, H., … Kenet, T. (2018). Maturation trajectories of cortical resting-state networks depend on the mediating frequency band. Neuroimage, 174, 57–68. doi:10.1016/j.neuroimage.2018.02.018

Khan, S., Michmizos, K., Tommerdahl, M., Ganesan, S., Kitzbichler, M. G., Zetino, M., … Kenet, T. (2015). Somatosensory cortex functional connectivity abnormalities in autism show opposite trends, depending on direction and spatial scale. Brain, 138(Pt 5), 1394–1409.

Klimesch, W. (2012). α-band oscillations, attention, and controlled access to stored information. Trends Cogn Sci, 16(12), 606–617. doi:10.1016/j.tics.2012.10.007

Klimesch, W., Doppelmayr, M., Schwaiger, J., Auinger, P., & Winkler, T. (1999). ’Paradoxical’ alpha synchronization in a memory task. Brain Res Cogn Brain Res, 7(4), 493–501. doi:10.1016/s0926-6410(98)00056-1

Koo, T. K., & Li, M. Y. (2017). A Guideline of Selecting and Reporting Intraclass Correlation Coefficients for Reliability Research (vol 15, pg 155, 2016). Journal of Chiropractic Medicine, 16(4), 346–346. doi:10.1016/j.jcm.2017.10.001

Lendner, J. D., Helfrich, R. F., Mander, B. A., Romundstad, L., Lin, J. J., Walker, M. P., Knight, R. T. (2020). An electrophysiological marker of arousal level in humans. Elife, 9. doi:10.7554/eLife.55092

Logothetis, N. K., Pauls, J., Augath, M., Trinath, T., & Oeltermann, A. (2001). Neurophysiological investigation of the basis of the fMRI signal. Nature, 412(6843), 150–157. doi:10.1038/35084005

Mann, E. O., & Mody, I. (2010). Control of hippocampal gamma oscillation frequency by tonic inhibition and excitation of interneurons. Nat Neurosci, 13(2), 205–U290. doi:10.1038/nn.2464

Manning, J. R., Jacobs, J., Fried, I., & Kahana, M. J. (2009). Broadband shifts in local field potential power spectra are correlated with single-neuron spiking in humans. J Neurosci, 29(43), 13613–13620. doi:10.1523/jneurosci.2041-09.2009

Manyukhina, V. O., Orekhova, E. V., Prokofyev, A. O., Obukhova, T. S., & Stroganova, T. A. (2022). Altered visual cortex excitability in premenstrual dysphoric disorder: Evidence from magnetoencephalographic gamma oscillations and perceptual suppression. PLoS One, 17(12), e0279868. doi:10.1371/journal.pone.0279868

Manyukhina, V. O., Prokofyev, A. O., Galuta, A. I., Goiaeva, D. E., Obukhova, T. S., Schneiderman, J. F., … Orekhova, E. V. (2022). Globally elevated excitation– inhibition ratio in children with autism spectrum disorder and below-average intelligence. Molecular Autism, 13. doi:10.1186/s13229-022-00498-2

Manyukhina, V. O., Rostovtseva, E. N., Prokofyev, A. O., Obukhova, T. S., Schneiderman, J. F., Stroganova, T. A., & Orekhova, E. V. (2021). Visual gamma oscillations predict sensory sensitivity in females as they do in males. Sci Rep, 11(1), 12013. doi:10.1038/s41598-021-91381-2

Maschke, C., Duclos, C., Owen, A. M., Jerbi, K., & Blain-Moraes, S. (2023). Aperiodic brain activity and response to anesthesia vary in disorders of consciousness. Neuroimage, 275, 120154. doi:10.1016/j.neuroimage.2023.120154

Mayhew, S. D., Coleman, S. C., Mullinger, K. J., & Can, C. (2022). Across the adult lifespan the ipsilateral sensorimotor cortex negative BOLD response exhibits decreases in magnitude and spatial extent suggesting declining inhibitory control. Neuroimage, 253, 119081. doi:10.1016/j.neuroimage.2022.119081

Mierau, A., Klimesch, W., & Lefebvre, J. (2017). State-dependent alpha peak frequency shifts: Experimental evidence, potential mechanisms and functional implications. Neuroscience, 360, 146–154. doi:10.1016/j.neuroscience.2017.07.037

Molina, J. L., Voytek, B., Thomas, M. L., Joshi, Y. B., Bhakta, S. G., Talledo, J. A., … Light, G. A. (2020). Memantine Effects on Electroencephalographic Measures of Putative Excitatory/Inhibitory Balance in Schizophrenia. Biol Psychiatry Cogn Neurosci Neuroimaging, 5(6), 562–568. doi:10.1016/j.bpsc.2020.02.004

Mullinger, K. J., Cherukara, M. T., Buxton, R. B., Francis, S. T., & Mayhew, S. D. (2017). Post-stimulus fMRI and EEG responses: Evidence for a neuronal origin hypothesised to be inhibitory. Neuroimage, 157, 388–399. doi:10.1016/j.neuroimage.2017.06.020

Mullinger, K. J., Mayhew, S. D., Bagshaw, A. P., Bowtell, R., & Francis, S. T. (2013). Poststimulus undershoots in cerebral blood flow and BOLD fMRI responses are modulated by poststimulus neuronal activity. Proc Natl Acad Sci U S A, 110(33), 13636–13641. doi:10.1073/pnas.1221287110

Murray, S. O., Kolodny, T., Schallmo, M. P., Gerdts, J., & Bernier, R. A. (2020). Late fMRI Response Components Are Altered in Autism Spectrum Disorder. Front Hum Neurosci, 14, 241. doi:10.3389/fnhum.2020.00241

Muthukumaraswamy, S. D. (2013). High-frequency brain activity and muscle artifacts in MEG/EEG: a review and recommendations. Front Hum Neurosci, 7. doi:ARTN 138 10.3389/fnhum.2013.00138

Muthukumaraswamy, S. D., & Liley, D. T. (2018). 1/f electrophysiological spectra in resting and drug-induced states can be explained by the dynamics of multiple oscillatory relaxation processes. Neuroimage, 179, 582–595. doi:10.1016/j.neuroimage.2018.06.068

Noonan, M. P., Crittenden, B. M., Jensen, O., & Stokes, M. G. (2018). Selective inhibition of distracting input. Behav Brain Res, 355, 36–47. doi:10.1016/j.bbr.2017.10.010

Orekhova, E. V., Butorina, A. V., Sysoeva, O. V., Prokofyev, A. O., Nikolaeva, A. Y., & Stroganova, T. A. (2015). Frequency of gamma oscillations in humans is modulated by velocity of visual motion. J Neurophysiol, 114(1), 244–255. doi:10.1152/jn.00232.2015

Orekhova, E. V., Manyukhina, V. O., Galuta, I. A., Prokofyev, A. O., Goiaeva, D. E., Obukhova, T. S., … Stroganova, T. A. (2023). Gamma oscillations point to the role of primary visual cortex in atypical motion processing in autism. PLoS One, 18(2), e0281531. doi:10.1371/journal.pone.0281531

Orekhova, E. V., Rostovtseva, E. N., Manyukhina, V. O., Prokofiev, A. O., Obukhova, T. S., Nikolaeva, A. Y., … Stroganova, T. A. (2020). Spatial suppression in visual motion perception is driven by inhibition: Evidence from MEG gamma oscillations. Neuroimage, 213, 116753. doi:10.1016/j.neuroimage.2020.116753

Orekhova, E. V., Stroganova, T. A., Schneiderman, J. F., Lundstrom, S., Riaz, B., Sarovic, D., … Hadjikhani, N. (2019). Neural gain control measured through cortical gamma oscillations is associated with sensory sensitivity. Hum Brain Mapp, 40(5), 1583–1593. doi:10.1002/hbm.24469

Orekhova, E. V., Sysoeva, O. V., Schneiderman, J. F., Lundström, S., Galuta, I. A., Goiaeva, D. E., … Stroganova, T. A. (2018). Input-dependent modulation of MEG gamma oscillations reflects gain control in the visual cortex. Sci Rep, 8(1), 8451. doi:10.1038/s41598-018-26779-6

Ossandón, J. P., Stange, L., Gudi-Mindermann, H., Rimmele, J. M., Sourav, S., Bottari, D., … Röder, B. (2023). The development of oscillatory and aperiodic resting state activity is linked to a sensitive period in humans. Neuroimage, 275, 120171. doi:10.1016/j.neuroimage.2023.120171

Ostlund, B. D., Alperin, B. R., Drew, T., & Karalunas, S. L. (2021). Behavioral and cognitive correlates of the aperiodic (1/f-like) exponent of the EEG power spectrum in adolescents with and without ADHD. Dev Cogn Neurosci, 48, 100931. doi:10.1016/j.dcn.2021.100931

Ouyang, G., Hildebrandt, A., Schmitz, F., & Herrmann, C. S. (2020). Decomposing alpha and 1/f brain activities reveals their differential associations with cognitive processing speed. Neuroimage, 205, 116304. doi:10.1016/j.neuroimage.2019.116304

Payne, L., Guillory, S., & Sekuler, R. (2013). Attention-modulated alpha-band oscillations protect against intrusion of irrelevant information. J Cogn Neurosci, 25(9), 1463–1476. doi:10.1162/jocn_a_00395

Peng, W., Babiloni, C., Mao, Y., & Hu, Y. (2015). Subjective pain perception mediated by alpha rhythms. Biol Psychol, 109, 141–150. doi:10.1016/j.biopsycho.2015.05.004

Perera, M. P. N., Mallawaarachchi, S., Bailey, N. W., Murphy, O. W., & Fitzgerald, P. B. (2023). Obsessive-compulsive disorder (OCD) is associated with increased electroencephalographic (EEG) delta and theta oscillatory power but reduced delta connectivity. J Psychiatr Res, 163, 310–317. doi:10.1016/j.jpsychires.2023.05.026

Peterson, E. J., & B., V. (2017). Alpha oscillations control cortical gain by modulating excitatoryinhibitory background activity. bioRxiv. doi:doi.org/10.1101/185074

Pietrelli, M., Samaha, J., & Postle, B. R. (2022). Spectral Distribution Dynamics across Different Attentional Priority States. J Neurosci, 42(19), 4026–4041. doi:10.1523/jneurosci.2318-21.2022

Podvalny, E., Noy, N., Harel, M., Bickel, S., Chechik, G., Schroeder, C. E., … Malach, R. (2015). A unifying principle underlying the extracellular field potential spectral responses in the human cortex. J Neurophysiol, 114(1), 505–519. doi:10.1152/jn.00943.2014

Razak, K. A., Binder, D. K., & Ethell, I. M. (2021). Neural Correlates of Auditory Hypersensitivity in Fragile X Syndrome. Front Psychiatry, 12, 720752. doi:10.3389/fpsyt.2021.720752

Riaz, B., Eskelin, J. J., Lundblad, L. C., Wallin, B. G., Karlsson, T., Starck, G., … Elam, M. (2022). Brain structural and functional correlates to defense-related inhibition of muscle sympathetic nerve activity in man. Sci Rep, 12(1), 1990. doi:10.1038/s41598-022-05910-8

Robertson, M. M., Furlong, S., Voytek, B., Donoghue, T., Boettiger, C. A., & Sheridan, M. A. (2019). EEG power spectral slope differs by ADHD status and stimulant medication exposure in early childhood. J Neurophysiol, 122(6), 2427–2437. doi:10.1152/jn.00388.2019

Roche, K. J., LeBlanc, J. J., Levin, A. R., O’Leary, H. M., Baczewski, L. M., & Nelson, C. A. (2019). Electroencephalographic spectral power as a marker of cortical function and disease severity in girls with Rett syndrome. J Neurodev Disord, 11(1), 15. doi:10.1186/s11689-019-9275-z

Sadaghiani, S., Uğurbil, K., & Uludağ, K. (2009). Neural activity-induced modulation of BOLD poststimulus undershoot independent of the positive signal. Magn Reson Imaging, 27(8), 1030–1038. doi:10.1016/j.mri.2009.04.003

Samaha, J., Iemi, L., Haegens, S., & Busch, N. A. (2020). Spontaneous Brain Oscillations and Perceptual Decision-Making. Trends Cogn Sci, 24(8), 639–653. doi:10.1016/j.tics.2020.05.004

Sheehan, T. C., Sreekumar, V., Inati, S. K., & Zaghloul, K. A. (2018). Signal Complexity of Human Intracranial EEG Tracks Successful Associative-Memory Formation across Individuals. J Neurosci, 38(7), 1744–1755. doi:10.1523/jneurosci.2389-17.2017

Shmuel, A., Augath, M., Oeltermann, A., & Logothetis, N. K. (2006). Negative functional MRI response correlates with decreases in neuronal activity in monkey visual area V1. Nat Neurosci, 9(4), 569–577. doi:10.1038/nn1675

Sohal, V. S., & Rubenstein, J. L. R. (2019). Excitation-inhibition balance as a framework for investigating mechanisms in neuropsychiatric disorders. Mol Psychiatry, 24(9), 1248–1257. doi:10.1038/s41380-019-0426-0

Stevens, F. L., Hurley, R. A., & Taber, K. H. (2011). Anterior cingulate cortex: unique role in cognition and emotion. J Neuropsychiatry Clin Neurosci, 23(2), 121–125. doi:10.1176/jnp.23.2.jnp121

Stevenson, C. M., Brookes, M. J., & Morris, P. G. (2011). β-Band correlates of the fMRI BOLD response. Hum Brain Mapp, 32(2), 182–197. doi:10.1002/hbm.21016

Tait, L., Özkan, A., Szul, M. J., & Zhang, J. (2021). A systematic evaluation of source reconstruction of resting MEG of the human brain with a new high-resolution atlas: Performance, precision, and parcellation. Hum Brain Mapp, 42(14), 4685–4707.

Taulu, S., & Hari, R. (2009). Removal of magnetoencephalographic artifacts with temporal signal-space separation: demonstration with single-trial auditory-evoked responses. Hum Brain Mapp, 30(5), 1524–1534. doi:10.1002/hbm.20627

van Kerkoerle, T., Self, M. W., Dagnino, B., Gariel-Mathis, M. A., Poort, J., van der Togt, C., & Roelfsema, P. R. (2014). Alpha and gamma oscillations characterize feedback and feedforward processing in monkey visual cortex. Proc Natl Acad Sci U S A, 111(40), 14332–14341. doi:10.1073/pnas.1402773111

Veerakumar, A., Tiruvadi, V., Howell, B., Waters, A. C., Crowell, A. L., Voytek, B., … Mayberg, H. S. (2019). Field potential 1/f activity in the subcallosal cingulate region as a candidate signal for monitoring deep brain stimulation for treatment-resistant depression. J Neurophysiol, 122(3), 1023–1035. doi:10.1152/jn.00875.2018

Vierling-Claassen, D., Cardin, J. A., Moore, C. I., & Jones, S. R. (2010). Computational modeling of distinct neocortical oscillations driven by cell-type selective optogenetic drive: separable resonant circuits controlled by low-threshold spiking and fast-spiking interneurons. Front Hum Neurosci, 4, 198. doi:10.3389/fnhum.2010.00198

Voytek, B., Kramer, M. A., Case, J., Lepage, K. Q., Tempesta, Z. R., Knight, R. T., & Gazzaley, A. (2015). Age-Related Changes in 1/f Neural Electrophysiological Noise. J Neurosci, 35(38), 13257–13265. doi:10.1523/jneurosci.2332-14.2015

Walker, M. C., & Semyanov, A. (2008). Regulation of excitability by extrasynaptic GABA(A) receptors. Results Probl Cell Differ, 44, 29–48. doi:10.1007/400_2007_030

Wianda, E., & Ross, B. (2019). The roles of alpha oscillation in working memory retention. Brain Behav, 9(4), e01263. doi:10.1002/brb3.1263

Wiest, C., Morgante, F., Torrecillos, F., Pogosyan, A., He, S., Baig, F., Tan, H. (2023). Subthalamic Nucleus Stimulation-Induced Local Field Potential Changes in Dystonia. Mov Disord, 38(3), 423–434. doi:10.1002/mds.29302

Wilkinson, C. L., & Nelson, C. A. (2021). Increased aperiodic gamma power in young boys with Fragile X Syndrome is associated with better language ability. Mol Autism, 12(1), 17. doi:10.1186/s13229-021-00425-x

Wilson, T. W., Slason, E., Asherin, R., Kronberg, E., Reite, M. L., Teale, P. D., & Rojas, D. C. (2010). An extended motor network generates beta and gamma oscillatory perturbations during development. Brain Cogn, 73(2), 75–84. doi:10.1016/j.bandc.2010.03.001

Zhang, F., Wang, F., Yue, L., Zhang, H., Peng, W., & Hu, L. (2019). Cross-Species Investigation on Resting State Electroencephalogram. Brain Topogr, 32(5), 808–824. doi:10.1007/s10548-019-00723-x

Zhang, Q., Cramer, S. R., Turner, K. L., Neuberger, T., Drew, P. J., & Zhang, N. (2023). High-frequency neuronal signal better explains multi-phase BOLD response. Neuroimage, 268, 119887. doi:10.1016/j.neuroimage.2023.119887

