## Supplementary materials for "Changes in high-frequency aperiodic 1/f slope and periodic activity reflect post-stimulus functional inhibition in the visual cortex"

**Running title: 'Post-stimulus inhibition in the visual cortex'**

Viktoriya O. Manyukhina[1,2], Andrey O. Prokofyev[1], Tatiana S. Obukhova[1], Tatiana A. Stroganova[1], Elena V. Orekhova[1]*

[1] Center for Neurocognitive Research (MEG Center), Moscow State University of Psychology and Education, Moscow, Russian Federation

[2] National Research University Higher School of Economics, Moscow, Russian Federation

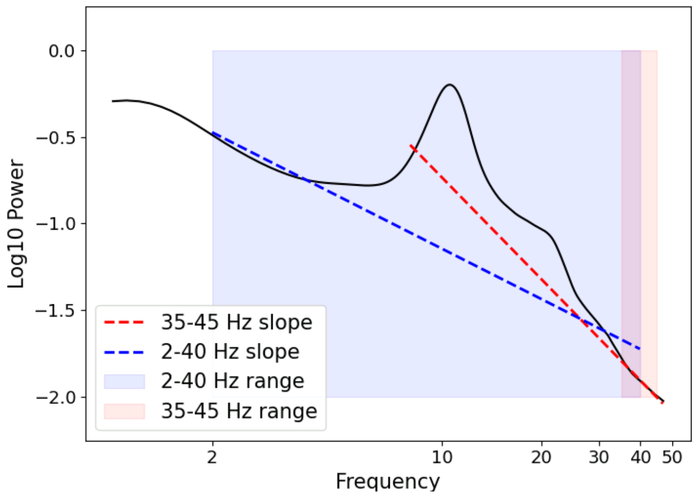


**Figure S1.** Illustration of aperiodic spectral slopes estimated for the group-averaged resting state condition using a linear approximation of the log-log spectrum in the 35-45 Hz range (red dashed line) or FOOOF in the 2-40 Hz range (blue dashed line).

### Results of the FOOOF analysis with aperiodic 'knee' mode.

To approximate the 'bend' in the aperiodic component we attempted to fit the power spectrum averaged over the posterior selection of gradiometers with FOOOF using a 'knee' mode.

Despite a good fit ($R^2$>0.93 for all subjects and conditions), there was high variability in the knee frequency between subjects: it ranged from 1.8 to 26.9 Hz (Fig. S2). This result contrasts with that of Ibarra Chaoul et al {Ibarra Chaoul, 2021 #49} who applied IRASA to separate aperiodic and periodic MEG PSD and found that a knee at 15 Hz was appropriate across all subjects and cortical regions.

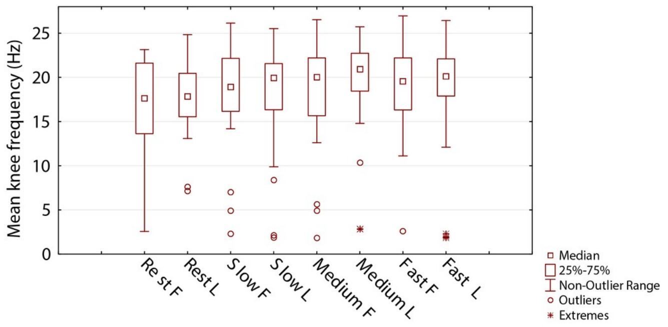


**Figure S2.** Variability of knee frequencies obtained using FOOOF with a 'knee' mode in the two phases of menstrual cycle (F – follicular and L – luteal) and two different conditions: rest and post-stimulus intervals (three types: 'after-slow', 'after-medium', and 'after-fast' preceding visual stimulation)

A comparison of the knee frequency in the same participants between two recording sessions has shown that the difference occasionally exceeded 5 Hz (Fig. S3).

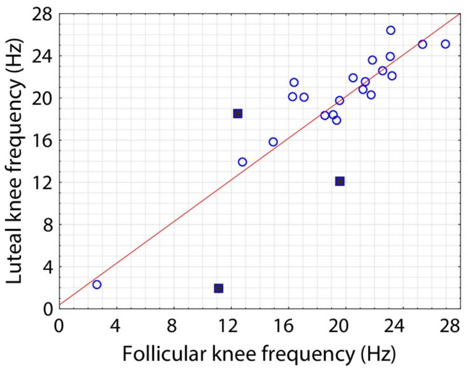


**Figure S3.** The correlation between knee frequencies estimated between two visits in the 'after-fast' condition. Squares mark cases with a large discrepancy between the estimated values.

Variability of the slope coefficients defined using FOOOF with the 'knee' mode was also high: it ranged from -0.35 to -7.30 in different subjects and conditions (Fig. S4). The very low slope (high exponent) values are at odds with those previously reported for MEG {Muthukumaraswamy, 2018 #46} and are strikingly different from those obtained in our study using a linear approximation of the PSD in the 35-45 Hz range (Fig. S5)

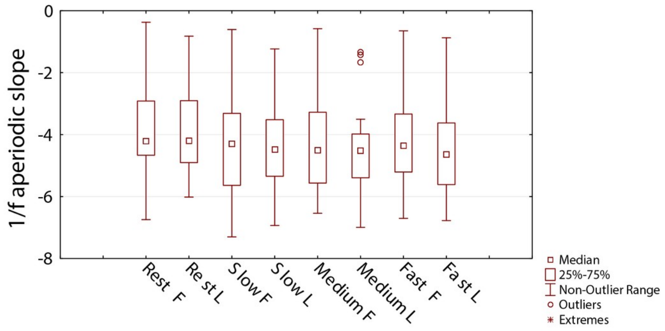


**Figure S4.** Variability of the aperiodic slope coefficients obtained using FOOOF with a 'knee' mode in the two phases of menstrual cycle (F – follicular and L – luteal) and two different conditions: rest and post-stimulus intervals (three types: 'after-slow', 'after-medium', and 'after-fast' preceding visual stimulation)

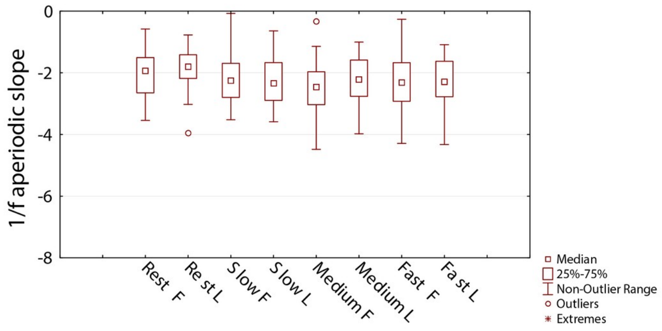

**Figure S5.** Variability of the aperiodic slope coefficients (in 35-45 Hz range) obtained using linear approximation of the PSD in log-log scale in the two phases of menstrual cycle (F – follicular and L – luteal) and two different conditions: rest and post-stimulus intervals (three types: 'after-slow', 'after-medium', and 'after-fast' preceding visual stimulation).

Considering that both the aperiodic knee frequency and the slope values obtained with FOOOF 'knee' mode were markedly different from those obtained in other MEG studies using other methods (Muthukumaraswamy & Liley, 2018; Chaoul et al, 2021), we concluded that estimation of the slope through linear approximation of the PSD in 35-45 Hz range, where periodic activity was absent, should give more interpretable results.

### Results of the analysis of the 35-45 Hz aperiodic slope at the marginal sensors

To test if the condition- and intensity-related differences in the 35-45 Hz aperiodic slope at the occipital selection of gradiometers (Fig. S6, red squares) can be explained by differences in muscle artifacts, we repeated the analyses for the selection of marginal posterior gradiometers closest to the neck muscles (Fig. S6, green squares).

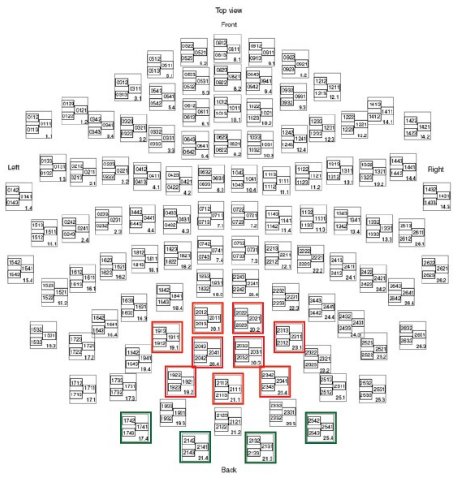


**Figure S6.** Sensors used for analysis in the 'sensor space'. Each colored square contains 3 sensors, of which only gradiometers (2 sensors in each location) were used for the analysis. Sensors marked in green were used to test the putative contribution of muscle artifacts to condition-related differences observed in the 35-45 Hz aperiodic slope, which were demonstrated in the main part of this paper for the selection of 9 pairs of gradiometers marked here in red.

For this selection, the rmANOVA with factors Condition (rest, post-stimulus) and Phase (follicular, luteal) revealed no significant effects of Condition ($F(1,24)=3.6$, $p=0.07$) or Condition x Phase interaction ($F(1,24)=0.01$, $p=0.90$).

Similarly, rmANOVA with factors Intensity ('after-slow', 'after-medium', 'after-fast') and Phase (follicular, luteal) revealed no significant effect of Intensity ($F(2,48)=1.55$, G-G epsilon = 0.96, $p=0.22$) or Intensity x Phase interaction ($F(2,48)=0.07$, G-G epsilon = 0.99, $p=0.93$).

The absence of significant effects of Condition and Intensity for the edge gradiometers indicates that the significant effects discussed in the main text which were found for the selection of 9 pairs of posterior gradiometers (Fig. S6, red squares) are unlikely to be explained by muscle artifacts.

### References


Ibarra Chaoul, A., & Siegel, M. (2021). Cortical correlation structure of aperiodic neuronal population activity. *Neuroimage, 245*, 118672. doi:10.1016/j.neuroimage.2021.118672

Muthukumaraswamy, S. D., & Liley, D. T. (2018). 1/f electrophysiological spectra in resting and drug-induced states can be explained by the dynamics of multiple oscillatory relaxation processes. *Neuroimage, 179*, 582-595. doi:10.1016/j.neuroimage.2018.06.068
